# Asymmetric Pendrin Homodimer Reveals its Molecular Mechanism as Anion Exchanger

**DOI:** 10.1101/2022.01.14.476289

**Authors:** Qianying Liu, Xiang Zhang, Hui Huang, Yuxin Chen, Fang Wang, Aihua Hao, Wuqiang Zhan, Qiyu Mao, Yuxia Hu, Lin Han, Yifang Sun, Meng Zhang, Zhimin Liu, Genglin Li, Weijia Zhang, Yilai Shu, Lei Sun, Zhenguo Chen

## Abstract

Pendrin SLC26A4 is an anion exchanger expressed in apical membranes of selected epithelia. Pendrin ablation causes Pendred syndrome, a genetic disorder disease associated with sensorineural hearing loss, hypothyroid goiter, and reduced blood pressure. However, its molecular structure has remained unknown limiting our understanding. Here, we determined the structures of mouse pendrin with symmetric and characteristically asymmetric homodimer conformations by cryo-electron microscopy. The asymmetric homodimer consists of an inward-facing protomer and an intermediate-state protomer, representing the coincident uptake and secretion process, and exhibits the unique state of pendrin as an electroneutral exchanger. This previously unrevealed conformation, together with other conformations we captured, provides an inverted alternate-access mechanism for anion exchange. Furthermore, our structural and functional data disclosed the properties of anion exchange cleft and interpreted the important pathogenetic mutations. These investigations shed light on the pendrin exchange mechanism and extend our structure-guided understanding of pathogenetic mutations.

**One-Sentence Summary:** Cryo-EM asymmetric Pendrin homodimer cracks its exchange mechanism and guides our understanding for pathogenetic mutations.

---

Pendrin encoded by the gene *SLC26A4* belongs to the solute carrier 26 (SLC26) family(*1*). It is a sodium-independent electroneutral Cl^−^/HCO_3_^−^ and Cl^−^/I^−^ exchanger expressed in the apical membrane of inner ear, thyroid and kidney epithelial cells (*2–6*) (Fig. 1A). In human, pendrin ablation cause the genetic disorder named Pendred syndrome(*7*), associated with sensorineural hearing loss, hypothyroid goiter, and reduced blood pressure(*6, 8*). Over 5% of the world’s population has hearing loss problem according to WHO data, among these about 3% is attributed to the hereditary *SLC26A4* pathogenetic variants(*9, 10*). Numerous genetics studies on the Pendred syndrome patients and various mouse models have provided insights about pendrin. In the inner ear, pendrin is found in the regions where endolymphatic fluid reabsorption occurs(*2*). Briefly, pendrin is expressed in epithelial cells of cochlear spiral prominence, endolymphatic sac, saccule, utricle and ampulla(*11*). The transport and exchange function of pendrin is believed to be critical in regulating the composition and pH stability of the inner ear’s endolymph. Pendrin knockout mice develops enlarged vestibular aqueduct (EVA) causing hearing impairment(*12*), and also affect acid-base balance of inner ear which will results in sensorineural hearing loss(*13, 14*). In kindney, pendrin localizes to type B and non-A, non-B intercalated cells (ICs) within the connecting tubule (CNT) and cortical collecting duct (CCD), where it mediates renal Cl^−^ absorption and HCO_3_^−^ secretion(*4, 6*). Within these apical plasma membranes, the abundance and activity of pendrin is significantly stimulated by angiotensin II and aldosterone, therefore would be beneficent to correct the alkalosis and maintain NaCl homeostasis to regulate blood pressure in kidney (*4, 15–17*). In the thyroid, pendrin is involved in I^−^ transport, however, its precise function is still under debate(*18*).

**Fig. 1.**
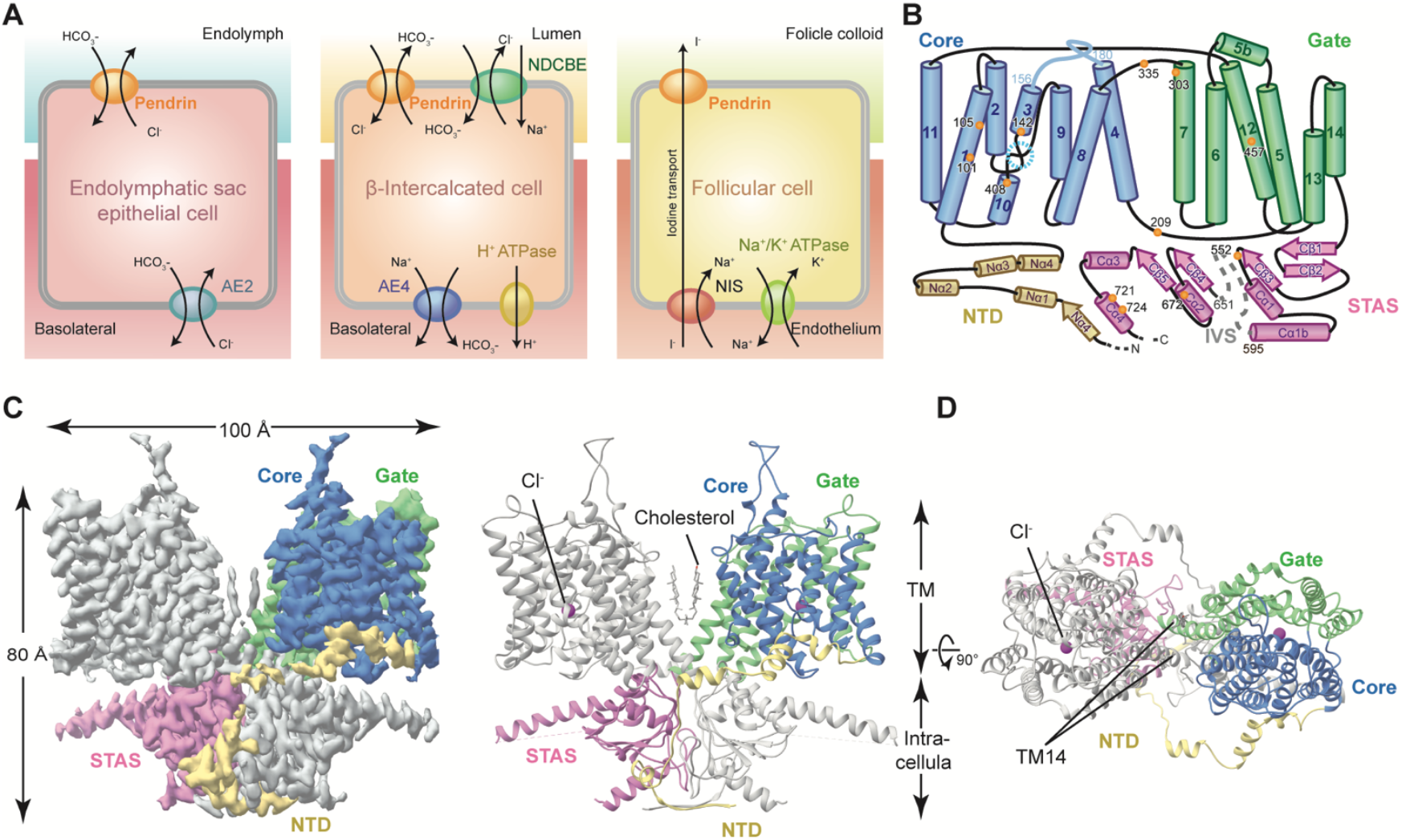
Cellular functions and cryo-EM structure of Pendrin. (**A**) Cellular functions of pendrin. In the inner ear **exchange** bicarbonate with chloride; in β-intercalated cells, it participates in urinary bicarbonate excretion with tubular chloride reabsorption; and in the thyroid, pendrin is involved in apical iodide transport. NIS, sodium iodide symporter; I^−^, iodide; Cl^−^, chloride; HCO_3_^−^, bicarbonate. (**B**) Topology of Pendrin. Some pathogenetic mutations in previous studies are shown as orange dots, the anion-binding pocket is marked by cyan circle. (**C**) Cryo-EM map and structural model of mouse pendrin-Cl^−^. One protomer is colored in grey, and the other is colored as (B). (**D**) View of (C) from outside the membrane. TM14 helix is marked.

The SLC26 members are normally anion transporters with diverse transport modes, accordingly they are divided into three general categories: the SO_4_^2-^ transporters SLC26A1 and SLC26A2; the Cl^−^/HCO_3_^−^ exchangers SLC26A3, SLC26A4 (pendrin) and SLC26A6; and the ion channels SLC26A7 and SLC26A9(*1, 19–21*). The human and dolphin SLC26A5 (prestin) was recently proved to be an electromotive signal amplifier, while its transport role remains unsolved(*22, 23*). Previously the mammalian SLC26A5 was reported to not as a transporter, although the invertebrate Slc26a5 does(*24, 25*). Among the whole SLC26 family, only SLC26A5(*22, 23*) and SLC26A9(*20, 21*) structures have been recently revealed by cryo-electron microscopy (cryo-EM), representing the electromotive signal amplifier and transporter structure, respectively.

Despite the importance of pendrin as an exchanger, its structure has remained unknown stumbling our understanding of the exchange mechanism. We investigated the structural and functional properties of mouse pendrin using single particle cryo-electron microscopy (Cryo-EM), electrophysiology and fluorescence microscopy. In the presence of one or two of the anions Cl^−^, HCO3^−^ or I^−^, we determined the structures of pendrin in three distinct states, including a symmetric inward-open dimer, a symmetric intermediate dimer and a characteristic asymmetric dimer composed of one inward-open protomer and another in the intermediate state. The discovered asymmetric homodimer architecture promotes us to understand the pendrin exchange toward an inverted alternate-access mechanism, with coincidently one protomer transporting in and another transporting out anions.

## Results

### Pendrin forms a symmetric homodimer in presence of Cl^−^

The full-length mouse pendrin with a N-terminal affinity tag were expressed in HEK293E cells and purified using glycol-diosgenin (GDN) in presence of 150 mM NaCl. The purified protein was concentrated to 1.7 mg/ml, and immediately used for cryo-EM grids freezing and subsequent data collection. The overall dimer structure was determined to 3.4 Å without symmetry, showing two protomers are almost identical. C2 symmetry was then applied for further refinement, which improved the reconstruction map to 3.3 Å resolution, allowing the unambiguous interpretation of the cryo-EM density map to atomic model. During protein purification and cryo-EM data processing, no other oligomers was observed, indicating pendrin forms a dimer as other family members(*20–23*) (fig. S1).

Pendrin purified in 150 mM NaCl (Pendrin-Cl^−^) forms a domain-swapped homodimer (Fig. 1, C and D). Each protomer is composed of a N-terminal domain (NTD) (residues 1-79), a transmembrane domain (TMD) (residues 80-514), a sulfate transporter and anti-sigma factor antagonist domain (STAS) (residues 535-735) containing the intervening sequence (IVS) (residues 596-650) and a C-terminal domain (CTD) (residues 736-780). Residues 1-17, 596-650 and 738-780 are missing in the model due to the flexibility. The NTD and STAS domains from each protomers interchange and form a dimeric knob at the cytoplasmic side of plasma membrane. On the other side, a spiky loop from each protomer sticks out from the membrane. The TMD follows the so-called UraA fold(*26, 27*), with 14 TMDs divided into two inverted repeats representing the core (helix TM 5-7 and 12-14) and gate regions (helix TM 1-4 and 8-11) (Fig. 1B, 2A). As with human prestin (PDB ID: 7LGU), significant cholesterol density was observed between the two TMDs (Fig. 1C-D).

### The anion-binding site between the core and gate regions within the TMD

In pendrin-Cl^−^ structure, an unambiguous Cl^−^ density is observed and defines the anion-binding site (Fig. 2, C and D). The Cl^−^ is embraced in the cavity between two short transmembrane helices TM3 and TM10, which is positioned at the apex of the inward-open intracellular vestibule (Fig. 2, F and G).

**Fig. 2.**
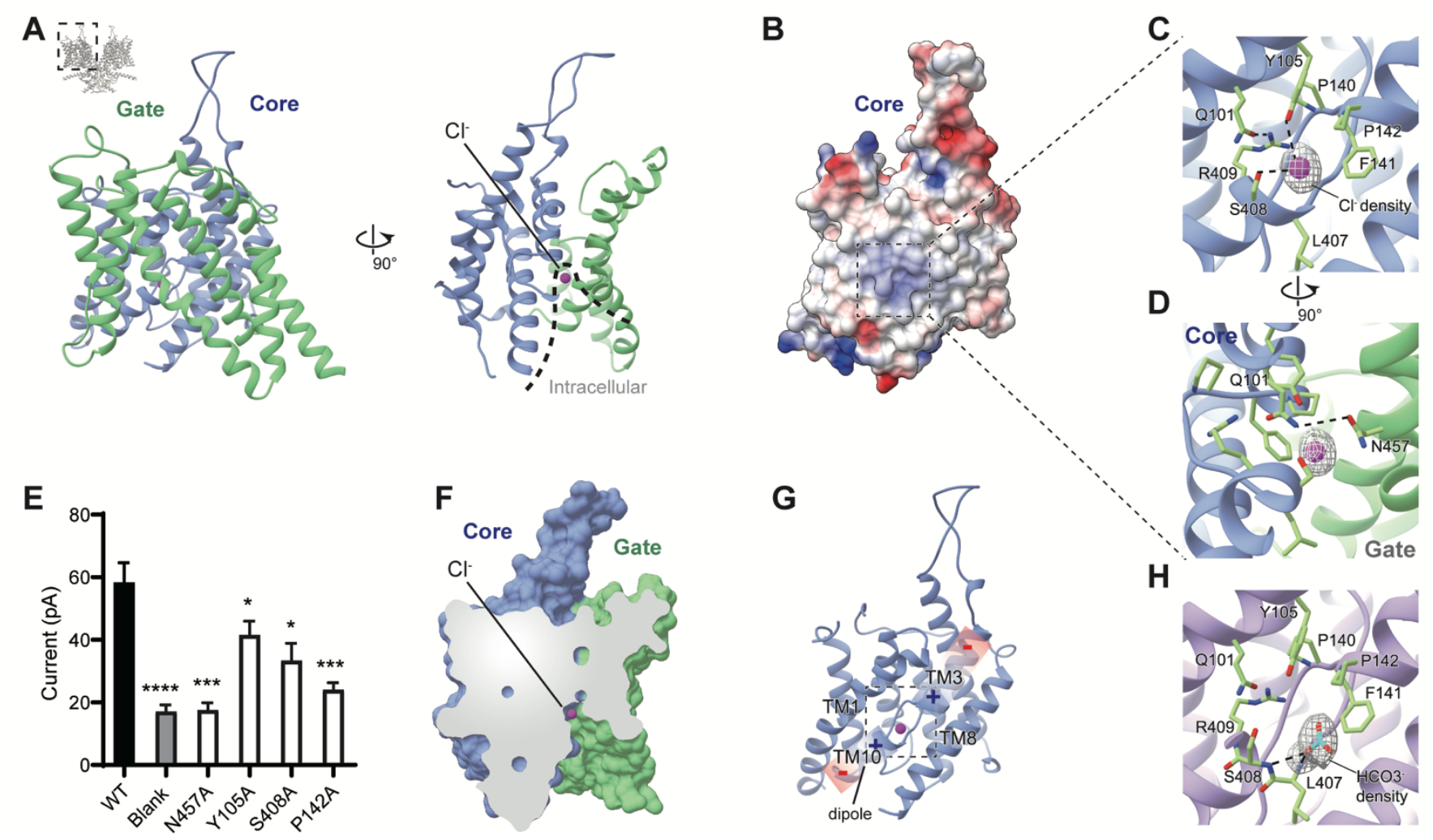
The TMD and anion binding pocket of pendrin. (**A**) Structural model of pendrin-Cl^−^ transmembrane domain (TMD) shown in ribbon representation. Core and gate regions are colored in lime green and dodger blue, respectively. Purple sphere indicates bounded Cl^−^, and dotted line describes the intracellular vestibule. (**B**) Electrostatic potential surface of the core region of pendrin-Cl^−^. (**C**) Details of the anion-binding pocket of pendrin-Cl^−^. Density representing Cl^−^ is shown in mesh mode. Interactions of Q101, Y105, L407 and S408 proximal to the pocket are shown. Hydrogen bond between Q101 and R409 is shown. (**D**) A 90° rotated view of (G) showing the hydrogen bond between Q101 and N457. (**E**) Currents of anion-binding site mutants (n = 9-19) at the voltage of 100 mV in HEK293T cells. Blank, non-transfected cells. ****P < 0.0001, ***P < 0.001 and *P < 0.05 versus the cells transfected with WT pendrin, unpaired t test. (**F**) Section of TMD. (**G**) Structural model of the core region of pendrin-Cl^−^. Gradient rectangles with charge symbol indicate the helical dipoles of TM3 and TM10. (**H**) Details of the anion-binding pocket of pendrin-HCO_3_^−^. Density representing HCO_3_^−^ is shown in mesh mode.

The Cl^−^ is mainly coordinated by the TM3-TM10 dipoles (Fig. 2G). S408 and Y105 directly interact with Cl^−^ by forming ionic bonds (Fig. 2C). The side chains of Y105 and Q101 form the partially positive charged binding pocket, which is quite common across the family(*20–23*). Additionally, the positively charged residue R409 indirectly stabilize Cl^−^ by interacting with the nearby residue Q101, which in turn interacts with S408. Mutation of R409 would alter anion binding and impair anion entrance and translocation(*28*), consequently R409H(*29*) leads to reduction of Cl^−^ and I^−^ transport. A406, L407 and three sequential hydrophobic residues P140, F141, P142 hallmark the partial hydrophobic feature of the pocket (Fig. 2B). Mutations of P140H(*29*) and P142R(*29*), though contributing positive charging, would cause the loss of anion transport or exchange activity, which may result from local structure disruption. Mutations of F141S(*28*) and P142R(*29*) were supposed to loss pi-pi stacking between F141 and P142 which will result into hearing problems. Additionally, residue N457 from the gate region also helps to stabilize the pocket by forming a hydrogen bond with Q101 (Fig. 2D).

Taken together, the local pocket architecture is crucial for anion binding and therefore important for pendrin’s anion transport and exchange function. We performed the electrophysiological recording and compared mutants P142A, Y105A, S408A and N457A with wild-type pendrin. Consistent with the structural findings, all four mutations significantly influenced the transport of Cl^−^ (Fig. 2E and fig. S1). Particularly, P142A and N457A showed severe reduction of Cl^−^ current, and both sites have pathogenetic mutations {P142R(*29*) and N457K(*28*)}. Both residues are supposed to be essential to the function of pendrin. Coincidently, the allelic residue of pendrin P142 is Alanine in prestin and SLC26A9. Therefore, P142 is assumed to have a specific role in pendrin to regulate Cl^−^ transport and exchange.

### HCO_3_^−^ binds the same pocket as Cl^−^

In order to obtain HCO_3_^−^-bound Pendrin (Pendrin-HCO_3_^−^), pendrin in Cl^−^ buffer was exchanged to a buffer containing HCO_3_^−^ using gel-filtration and frozen for cryo-EM data collection. The cryo-EM structure was determined to 3.5 Å resolution with C2 symmetry. Similar as pendrin-Cl^−^, pendrin-HCO_3_^−^ also forms a symmetric homodimer with two inward-open protomer. Anion HCO_3_^−^ was clearly observed as well in the binding pocket, which is about 2 Å away from Cl^−^ binding position (Fig. 2H). Though Y105 is too far to interact with, HCO_3_^−^ forms extra hydrogen bonds with the backbone nitrogen of S408 and L407.

### Pendrin forms asymmetric homodimer in presence of two different anions

As abovementioned, in the presence of a single anion, Cl^−^ or HCO_3_^−^, only inward-open pendrin was observed in our cryo-EM structures. The inward-open state may represent the most thermodynamically stable conformation but not dynamically favorite for anion transport and exchange. However, in physiological circumstance, pendrin is exposed a mixture of different anions. As an exchanger, pendrin could bind either anion (of the exchange pair) at both inward- and outward-open states. Therefore, we investigated pendrin in a buffer containing two different anions, such as Cl^−^/HCO_3_^−^, Cl^−^/I^−^, or HCO_3_^−^ /I^−^ anion pairs.

Pendrin purified in 100 mM NaCl was mixed with the same volume of 300mM NaHCO_3_, resulting a buffer containing 50 mM NaCl and 150 mM NaHCO_3_. The sample was then concentrated and used for cryo-EM data collection. The data was processed without symmetry. Surprisingly, the cryo-EM structural determination of pendrin in presence of Cl^−^/HCO_3_^−^ (pendrin-Cl^−^/HCO_3_^−^) mixture revealed three distinct states. Only ∼20% percent particles remained symmetric inward-open states. Another 20% percent particles represent a symmetric intermediate state in which both protomers adopt intermediate state, which is also observed in SLC26A9 and prestin structures (*21–23*). Interestingly, the majority of particles (∼60%) form asymmetric homodimers composed of one inward-open protomer and another intermediate protomer (Fig. 3A). Apparent anion densities were observed in symmetric inward-open state, which is assigned to Cl^−^ occupying identical position as that of pendrin-Cl symmetric structure. However, there is no clear anion density identified in the binding pocket of either symmetric intermediate state or asymmetric homodimer structure, which may due to the lower local resolution about 4 Å.

**Fig. 3.**
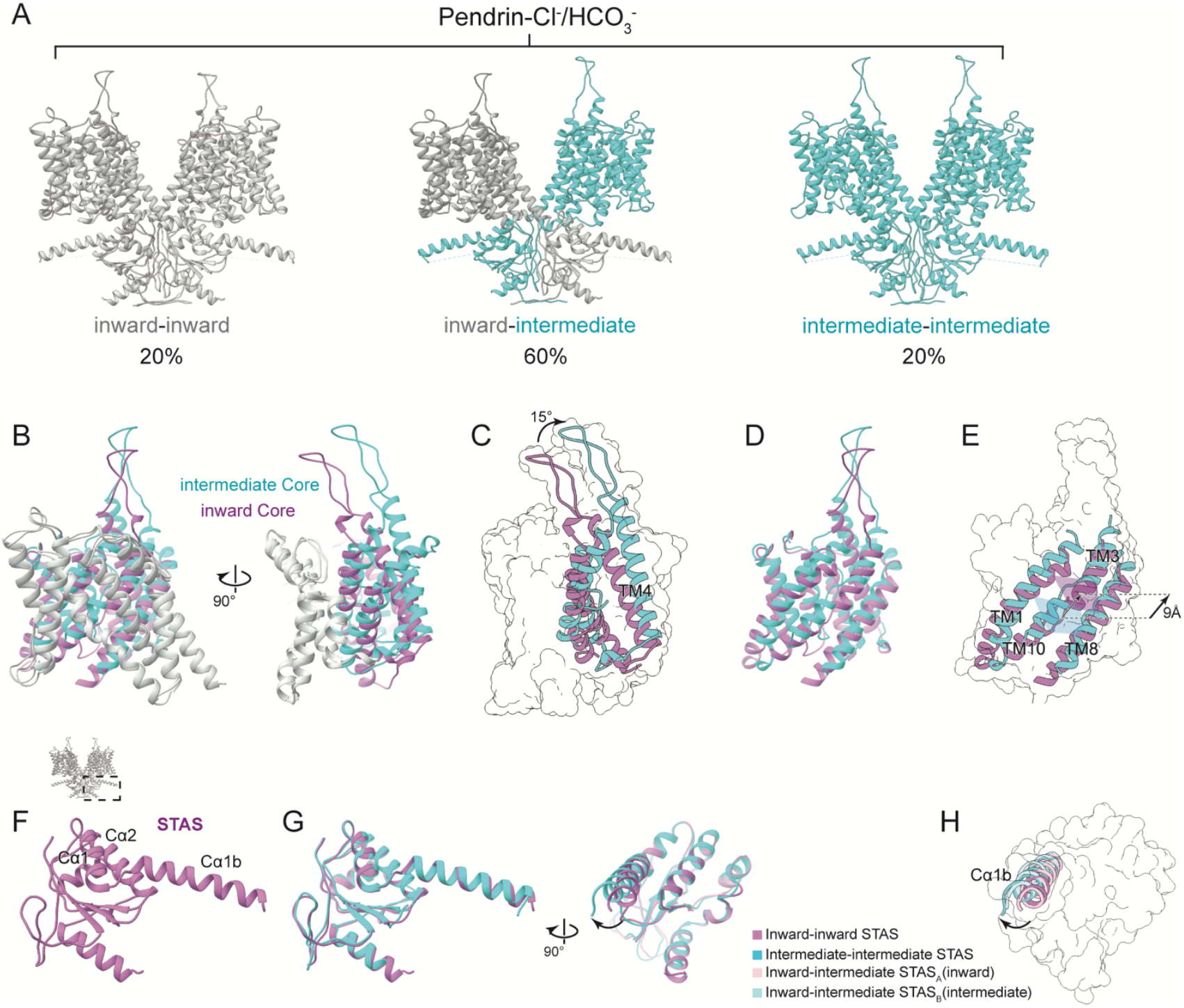
Comparison of inward-open and intermediate states of pendrin. (**A**) Three conformations of pendrin-Cl^−^/HCO_3_^−^. (**B**) Structural superimposition of the TMDs of inward (orchid and grey) state and intermediate state (cyan and grey) of pendrin. (**C**) Comparison of the core regions in the inward state and the intermediate state. TM4 of intermediate state rotates roughly by 15°. (**D**) The core region of (B). (**E**) As the core region rotates, the anion-binding pocket indicated by translucent tetragon translates 9 Å towards the extracellular side. (**F**) STAS domain of inward state of pendrin-Cl^−^/HCO_3_^−^. (**G**) Structural superimposition of the STAS domains of inward (orchid) state and intermediate state (cyan) (**H**) Differences of the helix Cα1b among four protomers. Inward-inward (orchid), intermediate-intermediate (cyan), inward-intermediate protomer A(pink), and inward-intermediate protomer B (light blue).

Similarly, the pendrin sample in the anion pair of Cl^−^/I^−^ (1:3) (pendrin-Cl^−^/ I^−^) was prepared and used for structure determination. Symmetric inward-open and asymmetric homodimers were observed with approximate percentage of 75% and 25%, respectively. Finally, the structures of pendrin in HCO3^−^/I^−^ (1:3) (pendrin-HCO3^−^/I^−^) was determined, which also revealed two distinct states: 75% symmetric inward-open and 25% asymmetric states. The combination of Cl^−^ and HCO_3_^−^ provides the most varieties of conformations, hereafter pendrin-Cl^−^/HCO3^−^ structures are used for illustration.

Remarkable changes among the TMD of the inward-open and intermediate pendrin protomer were observed. The major difference is the relative positions of the gate and the core regions (Fig. 3B). When the gate regions are superimposed, the core region in the intermediate state (versus the inward-open state) rotates roughly by 15° and translates 9 Å towards the extracellular side (Fig. 3, C to E). The movement is characteristic in SLC26 family members and the relative value is the biggest among published data(*21–23*), while the uppermost conformation would end up with the outward-open state of SLC4A1 (anion exchanger 1, AE1)(*27*). This outward-open state is believed to release the bound anion, and carry another anion of the exchange pair to reverse the process(*27*).

In addition, the density of relatively stable cholesterol can be seen in all conformations, despite the completeness differences. Since cholesterol is believed to influence the localization and diffusion of prestin in plasma membrane(*30*), this may be characteristic for the interactions between SLC26 family members and plasma membranes.

### The STAS domain modulates pendrin transport function

The STAS domain takes 4 β-strands as the core which is surrounded by 4 α-helixes, extending the lateral helix Cα1b to link IVS (Fig. 3F). The starting residues 515-545 of STAS domain form a long loop region, however, it is well resolved due to the interactions with Cβ3, Cα5 and NTD. The pathogenetic mutations we investigated with electrophysiology, Y530H, T721M, and D724N, which severely affected Cl^−^ transport, are located around this loop region (Fig. 4, B and C) (*31*). This indicate that the structural stability of the STAS domain has dramatic influence on the transport function. Similar to prestin and SLC26A9, pendrin’s dimerization interface is mainly formed by STAS domains. The STAS domains of two protomers contact closely face to face on a relatively flat surface, and two NTD’s Nβ1s parallel inversely below the STAS domains. In the dimerization interface, S552 forms a hydrogen bond with S666 from the other protomer, which would be destroyed by pathogenetic mutation S552I(*31*) (Fig. 4, E and F). Referring to the complete loss of Cl^−^ permeability in electrophysiological experiment, we hypothesized that the instability of dimerization would disable the transport function of Pendrin (Fig. 4D).

**Fig. 4.**
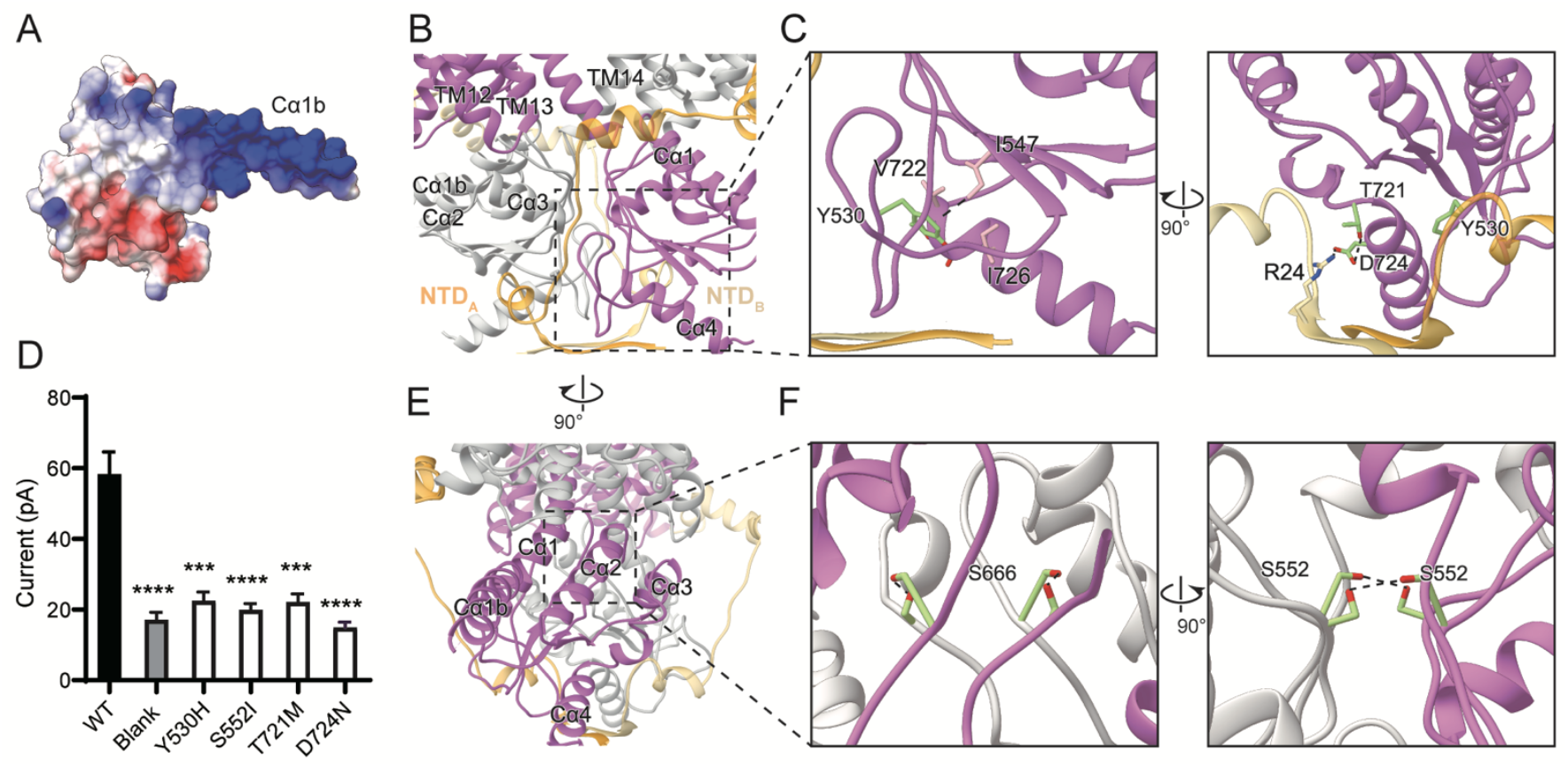
STAS domain of pendrin. (**A**) Electrostatic potential surface of the STAS domain of inward state of pendrin-Cl^−^/HCO_3_^−^. (**B**) The dimer interface of inward-inward state of pendrin-Cl^−^/HCO_3_^−^. Protomer A is colored in orchid and orange, protomer B is colored in grey and khaki. (**C**) Details of STAS domain. Y530, T721, D724 and their interactions are indicated by dot lines. (**D**) Currents of pathogenetic mutants on STAS domain in HEK293T cells analyzed at the voltage of 100 mV. n = 10-19. Blank, non-transfected cells. ****P < 0.0001 and ***P < 0.001 versus the cells transfected with WT pendrin, unpaired t test. (**E**) A 90° rotated view of (B). (**F**) Details of interface between two STAS domains. Hydrogen bonds between S552 and S666 are indicated.

The STAS of one protomer not only interacts with the STAS of another protomer, but also contacts the TMD of another protomer, as the platform formed by Cα1, Cα1b, and Cα2 is directly below the anion transport pathway (Fig. 4B). Moreover, an anion pre-binding site between loop Cβ3-Cα1 and loop Cβ4-Cα2, according to the rat prestin X-Ray structure (*32*), forms an interface with TM12, TM13 and TM14. Pathogenetic mutations Y556C, F667C, and G672E of this region, would significantly affect Cl^−^ and I^−^ transport(*29*). When the TMDs were superimposed, the helix Cα1b rotated about 6° from inward state to intermediate state (Fig. 3, G and H). Interestingly, Cα1b provides a completely positively charged platform, therefore the movement may significantly contribute to the initiation of anion transport within STAS domain, subsequently facilitating the anion transport or exchange (Fig. 4A).

### Cl^−^/HCO_3_^−^ and Cl^−^/I^−^ exchange function of pendrin

To verify the Cl^−^/HCO_3_^−^ exchange function of pendrin, we used pH sensitive fluorescent probe BCECF to reflect HEK293T intracellular concentration of HCO_3_^−^ in different bath(*33*). And for Cl^−^/I^−^ exchange function, halides-sensitive EYFP with different sensitivity to Cl^−^ and I^−^ was co-transfected with pendrin to detect the intracellular changes of Cl^−^ and I^−^ concentration(*34*). As a positive control, the wild type pendrin showed remarkable exchange functions in both experiments (Fig. 5, H and M). Thereafter, pathogenetic mutations reported to affect the exchanger function (E303Q, F335L, G209V and G672E) were tested and distinct responses were detected (Fig. 5A).

E303 is located on the core-gate interface, although it is far from the anion pathway, mutation E303Q still loose anion exchange capability (Fig. 5E). According to both exchange experiments, E303Q is no longer permeable to Cl^−^, resulting in the blocked exchange, but still retains the permeability to I^−^ (Fig. 5, F, G, K and N). Glutamine would reverse the nearby surface charge (from negative to positive), thus affecting the thermodynamics of conformational change. F335 is located at the protein-lipid interface and appears to have strong interactions, as excess lipid density is seen next to F335 in cryo-EM map (Fig. 5D). In fluorescence experiments, F335L was substantially weakened on both exchanges (Fig. 5, F, G, L and O). Structurally, the side chain of leucine may disrupt protein-lipid interactions, thereby impairing the allosteria of TMD. It may indicate that relatively immobilized protein-lipids interactions are essential for the normal function of pendrin. G209 is located on the interface between TMD and STAS domain, and pendrin G209V was reported to located on plasma membrane (PM) and show severe reduction of I^−^ transport (Fig. 5B)(*28*). However, our fluorescence assays not only demonstrated the functional impairment but also displayed intracellular localization of pendrin G209V (Fig. 5, F and I). According to the cryo-EM structure, side chain of valine would increase steric hindrance of core-gate interface and contribute the positive surface charge. These changes might influence the allosteria of TMD and pendrin localization on plasma membrane. G672, conserved with Rat Prestin, is located at the hypothetic pre-binding site in the STAS domain (Fig. 5C). In Cl^−^/I^−^ exchange assays, G672E lost I^−^ transport capacity, but maintains Cl^−^ permeability (Fig. 5, F and J). The long side chain of glutamic acid would extend into the cleft, perhaps altering pendrin’s anion selectivity. In summary, the pendrin structures provide a plausible guideline for us to understand pathogenetic mutations at a molecular level.

**Fig. 5.**
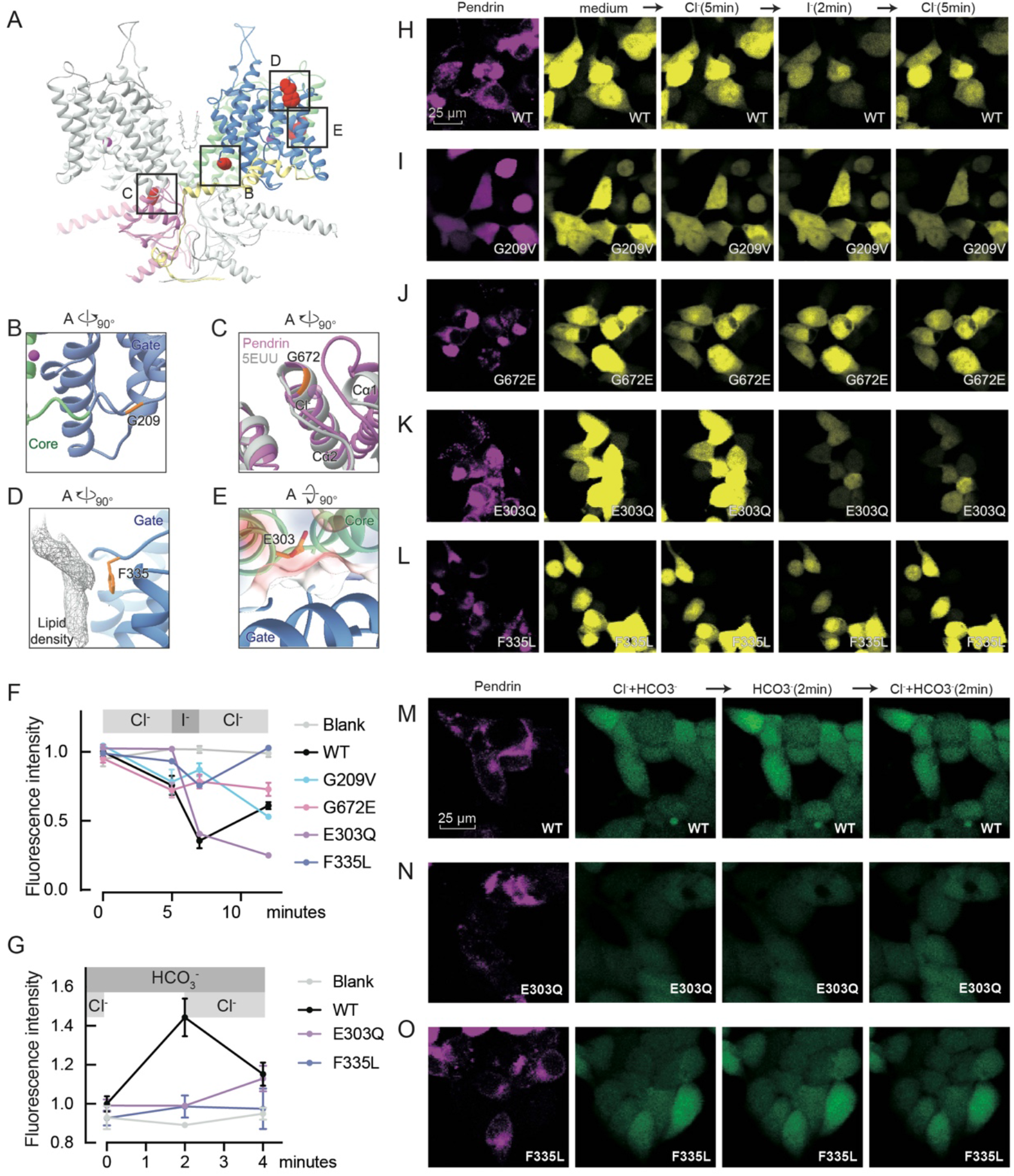
Pathogenetic mutations of PSD and fluorescence exchange assays. (**A**) Four pathogenetic mutations are indicated by sphere mode, G209, G672, E303 and E335. (**B**) Details of G209 location. (**C**) Superimposition of pendrin-Cl^−^ STAS domain(orchid) and rat prestin STAS domain (PDB ID: 5EUU, grey). (**D**) lipid density neighboring F335 is indicated in mesh mode. (**E**) Local electrostatic potential surface of E303. (**F**) Fluorescence intensity change in HEK293T cells in Cl^−^/I^−^ exchange assay. n = 5. Blank, EYFP-transfected cells. (**G**) Fluorescence intensity change in HEK293T cells in Cl^−^/HCO_3_^−^ exchange assay. n = 5. Blank, non-transfected cells. (**H-L**) Micrographs of pendrin+EYFP-transfected cells. Shooting parameters keep the same in each group. Note that G209V-pendrin is localized differently (I). (**M-O**) Micrographs of pendrin-transfected cells treated with BCECF. Shooting parameters keep the same in each group.

### Mapping of critical pathogenetic mutations

Pendrin is the best-studied family member due to the enormous mutations that have been implicated in Pendred syndrome. These mutations showed a very dense distribution present in almost every region of pendrin(*31*). Notably, a single missense would lead to dysfunction or loss of function. Our cryo-EM pendrin structures make it possible to rationalize previous mutational studies. Of all the identified mutations, we analyzed those with a well-defined cellular localization verified by experiments (table. S3). Therefore, we expect to explain how PM localization dysfunction is derived, and why membrane-localized pendrin mutants cannot transport anions properly.

There are six missense variants, which still secure pendrin PM localization, vary in function impairment. R409H located in the binding pocket changes the hydrogen bond that guarantees stability, causing reduction of Cl^−^ and I^−^ transport(*28*). Anion access pathway related mutation E303Q would reverse local charge to cause function impairment. Two mutation variants apart from anion exchange pathway, F335L would influence the protein-lipid interaction, and G209V could affect core-gate interface stability, therefore they could affect exchange function as well. The rest two S28R and C565Y variants remain unclear on function impairment interpretation.

Mostly, 14 out of the 22 missense variants would mis-localize in the endoplasmic reticulum (ER) or cytoplasm. Among these 7 variants, in TMD (G102R, P123S, V239D, E384G, N392Y, L445W, and G497S), introduce new steric hindrances between different TM helixes, which would alter the position and orientation of the helixes, thus eventually disordering the overall arrangement of core and gate regions. Besides, V138 is closely connected to the binding pocket, and mutation V138F will affect organization of the loop next to TM3. Similarly, T410M providing a large side chain will also affect the loop next to TM3. For R185T at the core-gate interface and L236P at the protein-lipids interface, the charge change may lead to local misfolding. It is noteworthy that mutations in intracellular STAS domain also affect pendrin’s PM localization. Y530H/S, T721M and H723R are concentrated at the bottom of pendrin, participating in the stabilization of STAS domain and NTD, and also determining the orientation of Cα4. Additionally, Y556C and G672E in TMD-STAS interface would only partially affect PM localization. This indicates that these mutations on the interface modify local stability which subsequently impair PM localization.

### Structural comparison of pendrin, prestin and SLC26A9

The overall fold of pendrin is similar to prestin(*22, 23*) and SLC26A9(*20, 21*), however, the atomic cryo-EM structures of pendrin reveal its intrinsic features of asymmetric homodimer as an exchanger, given the symmetric homodimers resolved for all other SLC26 family members(*20–23*). The structures of mouse SLC26A9 have two states: inward-open (PDBID: 6RTC) and intermediate (PDBID: 6RTF) states(*21*). The dolphin prestin structures include several states, including an inward-like prestin-SO_4_^2−^ (PDBID: 7S9C), intermediate-like prestin-Cl^−^ (PDBID: 7S8X) and several other states in between these two states(*23*).

When superimposing the inward-state protomers, the TMD regions of pendrin, prestin and SLC26A9 are relatively similar (RMSD<1.3Å, table. S2). The major difference is that the STAS domain of SLC26A9 has a distinct angular offset from pendrin and prestin, while the latter two basically overlap with each other.

The TMDs of pendrin, prestin and SLC26A9 in the intermediate state show larger differences than the inward states. Among the intermediate state protomers, pendrin’s core region has the largest rotation, surpassing that of mouse SLC26A9. Therefore, the anion binding pocket of the intermediate state pendrin is also the uppermost and close to the outward-open state of AE1 (PDBID: 4YZF, Fig 6A). When focused on anion binding pocket, within a conserved framework, residue differences were found at key sites between pendrin, prestin and SLC26A9. Sequence alignment showed that pendrin residues Q101, F141, L407 and S408 are invariant, but not Y105, P140, P142 and R409 (fig. S10). Different from the allelic residue of prestin and SLC26A9’s phenylalanine, pendrin Y105 has the phenolic hydroxyl group, which increases the charge of binding pocket and enhances the electrostatic attraction to anions. Pendrin’s P140-F141-P142 hydrophobic fragment has slightly different allelic residues in prestin and SLC26A9, PFA and TFA, respectively. The diversity of these three consecutively amino acids tunes the surface shape and charge of the pocket, and may cause differences in interaction intensity between the pocket and anions. Finally, pendrin R409 is conserved in most SLC26A members except SLC26A1, SLC26A2 and SLC26A9. In SLC26A1 and SLC26A2 it is replaced by Lysine, which might be related to the specific function in SO_4_^2−^ transport; while in SLC26A9, this residue was replaced by Valine (fig. S9). Within the binding pocket, pendrin R409 provides the only positive charge and forms multiple hydrogen bonds with neighboring residues for stability, as does in Prestin. However, Valine in SLC26A9 only has a hydrophobic short side chain, which weakens the overall positive charge of pocket. In summary, surface charge of the binding pocket may directly define the anion selectivity, which may be the reason why these three members are so different in terms of anion selectivity and function.

**Fig. 6.**
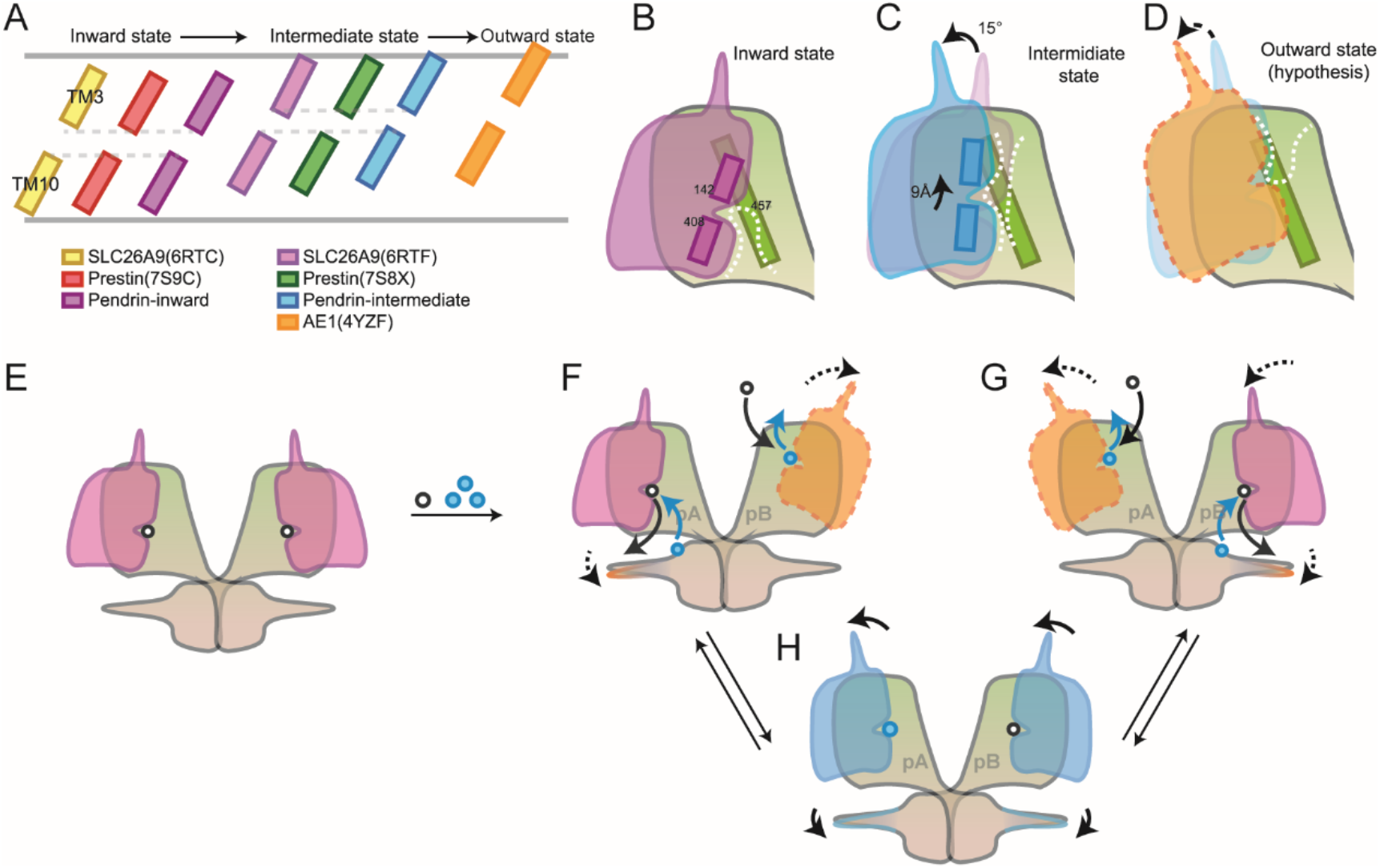
Comparison among family members and exchange mechanism. (**A**) TM3 and TM10 defined the pocket position of representative states of SLC members pocket. (**B-D**) Schematic representation showing a complete alternate opening in TMD. (**E-H**) Schematic representation showing a hypothetic exchange mechanism. (E) A thermodynamically stable symmetric inward homodimer in resting state. (F) binding of anions pair activates the transport of one protomer. (G) Allosteria of TMD and STAS from one protomer interact the transport of another protomer, which determines the inverted alternate-access mechanism. pA and pB: protomer A and protomer B, respectively.

## Discussion

The competitive binding of anion pair in binding pocket, which pivot points the exchange cleft’s access direction, determines the conformation of the TMD and probably modulates that of STAS as well. Here, we have resolved a wide spectrum of pendrin structures flash frozen under various conditions. On the basis of structural analysis together with physiological and biochemical assays, we hypothesized the working model of pendrin as an anion exchanger. The symmetric inward-open conformation most likely represents the energy favorite state, existing in the single binding-anion circumstance, Cl^−^ for instance (Fig. 6E). When bicarbonate was added to a high concentration, HCO_3_^−^ replaces Cl^−^ from the same binding site in protomer B. This causes local conformation changes, with the binding pocket of protomer B translating towards the extracellular side, while protomer A stays unchanged (Fig. 6F; movie. S1). To accommodate this change, the core region rotates about 15° against the gate region, indicating the elevator transport mechanism(*21*)(Fig. 6C). Eventually this conformational change would end up with the outward-open state(*27*) (Fig. 6D), at which HCO_3_^−^ diffuses out and Cl^−^ binds to reverse the exchange process (Fig. 6F). The significant rotation of Cα1b of protomer B, occurring at the end stage of anion releasing prior to anion uptake, may facilitate the on-set of protomer A’s anion secretion process (Fig. 6H). Therefore, the coincidence of secretion and uptake in this asymmetric homodimer, shapes the molecular basis of electroneutral exchange of pendrin with the so-called inverted alternate-access mechanism. To the best of our knowledge, it was never observed before for any other anion exchanger. Moreover, some of the important pathogenetic mutations were mapped on the structure, functional studies were also performed to interpret the structure-function relationships. All above-mentioned would provide a framework for us to understand more pathogenetic mutations and could shed light on therapeutic discovery.

## Acknowledgments

We thank Center of Cryo-Electron Microscopy, Fudan University for the supports on cryo-EM data collection.

## Funding

This work was supported by the National Natural Science Foundation of China (81900729 to L.S., 31970146 to Z.C.), National Key Research and Development Program of China grant 2020YFA0908201(Y.S.), National Natural Science Foundation of China grant 82171148(Y.S.), and the Science and Technology Commission of Shanghai Municipality 21S11905100 (Y.S).

## Author contributions

ZC, LS and YS conceived the project; QL cloned, expressed and purified proteins; XZ, ZC and QL collected and processed cryo-EM data. QL and LS built the atomic models; YC and FW performed electrophysiology assays; QL and HH performed fluorescence assays; all authors are involved to analyze and discuss the data; QL, LS and ZC wrote the manuscript and reviewed by all the authors.

## Competing interests

Authors declare that they have no competing interests.

## Data and materials availability

The cryo-EM density maps and coordinates have been deposited in the Electron Microscopy Data Bank (EMDB) and Protein Data Bank (PDB), respectively, with accession codes 32555 and 7WK1 for pendrin-Cl, 32561 and 7WK7 for pendrin-HCO_3_, 32577 and 7WL8 for pendrin-Cl/I_ii_, 32580 and 7WLB for pendrin-Cl/I_im_, 32576 and 7WL7 for pendrin-Cl/HCO_3ii_, 32578 and 7WL9 for pendrin-Cl/HCO_3im_, 32583 and 7WLE for pendrin-Cl/HCO_3mm_, 32574 and 7WL2 for pendrin-HCO_3_/I_ii_, 32579 and 7WLA for pendrin-HCO_3_/I_im_.

## Supplementary Materials

Materials and Methods

Figs. S1 to S10

Tables S1 to S4

References (35-46)

Movies S1

## Notes

### Competing Interest Statement

The authors have declared no competing interest.

