## Supplemental materials for "Asymmetric Pendrin Homodimer Reveals its Molecular Mechanism as Anion Exchanger"

**This PDF file includes:**

Materials and Methods

Figs. S1 to S10

Tables S1 to S4

References (35-46)

Movies S1. Conformational change of pendrin homodimer from the inward-open to intermediate state

### Materials and Methods

#### Constructs and cell culture

DNA encoding full-length mouse pendrin (uniport ID: Q9R155-1) was synthesized and subcloned into the pEZT BacMam vector, with a N-terminal 3X Flag tag and a 3C protease cleavage site. For fluorescence assay, full-length mouse pendrin was shuttled into a pcDNA 3.1 vector and N-terminal tags were replaced by a mCherry or EGFP fluorescent protein sequence. Single point mutations were PCR amplified and subcloned based on wild type plasmids.

Sf9 cells were cultured in ESF921 medium at 27 °C and used for BacMam virus amplification. P3 virus used for infection was precipitated by PEG 6000 and re suspended with 1/10 volume of OPM medium. HEK293E cells used for protein expression and purification were cultured in OPM medium at 37 °C and 5% CO2. Adherent HEK293T cells for fluorescence assay were cultured in DMEM medium supplemented with penicillin/streptomycin and 10% FBS.

#### Protein expression and purification

HEK293E cells cultured at 37 °C with a density of ~$2.3\times10^6$ /ml were infected with resuspended BacMam virus at a volume ratio of 100:1. A final concentration of 10 mM sodium butyrate was added 12 hours after infection, and cells were cultured at 37 °C for another 48 hours. For fluorescence assays, adherent HEK293T cells were cultured in 6-well plate with a cell density of ~70%, and transfected with plasmids-PEI mix. For Electrophysiological recoding, HEK293T cells were transiently transfected with plasmids using Lipofectamine 3000™ reagent (Invitrogen) according to the manufacturer's instructions. After transfection for 24 h, the GFP-positive cells were selected randomly for patch-clamp recording.

All subsequent purification procedures were carried out at 4 °C. Two liters of HEK293E cells were harvested and resuspended in buffer containing 20 mM Tris pH 8.0, 150 mM NaCl and protease inhibitors (0.8 mM aprotinin, 2 mg/ml leupeptin and 2 mM pepstatin A), and 1.5% (w/v) digitonin power was added into cell suspension and incubated for three hours. After centrifugation at 18000 RPM for an hour, the supernatant was filtered through a 0.45 μm filter and incubated with anti-flag affinity resin. The resin was washed with 20 mM Tris HCl pH 8.0, 150 mM NaCl, 5 Mm Mg-ATP, 0.02% (w/v) glycol-diosgenin (GDN) to remove heat shock proteins, and eluted with 0.3mg/mL Flag peptide.

For pendrin-Cl, the concentrate was loaded onto a size-exclusion chromatography (SEC) column (Superose 6 Increase 10/300 GL) equilibrated with a buffer containing 20 mM Tris HCl pH 8.0, 150 mM NaCl, 0.02% GDN (fig. S1G). The peak fraction was analyzed by SDS-PAGE (fig. S1H) and concentrated to ~1.7mg/mL for cryo-EM sample preparation. For pendrin-Cl/$HCO_3$ and pendrin-Cl/I, an SEC buffer containing 20 mM Tris pH 8.0, 100 mM NaCl, 0.02% GDN was used. And equal volume of 300 mM NaHCO3 (the final pH of sample pendrin- Cl/$HCO_3$ was 8.3) or NaI was added into peak fractions before concentrating. For pendrin pendrin-$HCO_3$ and pendrin-$HCO_3$/I, an SEC buffer containing 20 mM HEPES pH 8.0, 60 mM $Na_2$SO4, 0.02% GDN was used to replace $Cl^-$, and a final concentration of 10 mM NaHCO3 was added into peak fractions during the concentration and the final pH of sample pendrin-$HCO_3$ was 8.3. Sample pendrin-$HCO_3$/I was added with a final concentration of 33 mM NaI based on sample pendrin-$HCO_3$.

**Cryo-EM grid preparation and data collection**

Each pendrin sample was concentrated to about 1.7 mg/mL, and 3 μL of the protein was placed on glow-discharged grids (NiTi Au 400# R1.2/1.3), then the grid was blotted for 2.0 s with blotting force -2 and flash-frozen in liquid ethane with Vitrobot Mark IV (Thermo Fisher Scientific®).

Some pendrin cryo-EM datasets (including pendrin-Cl, pendrin-$HCO_3$ and pendrin-$HCO_3$/I) were collected on Titan Krios (Thermo Fisher Scientific®) operated at 300 kV, equipped with K2 Summit direct electron detection device (Gatan®) and BioQuantum energy filter (Gatan®) set to a slit width of 20 eV. Automated data acquisition was carried out with SerialEM software(*1*) through the beam-image shift method(*2*). Others (pendrin-Cl/$HCO_3$, pendrin-Cl/I) were collected on the same microscope with the camera upgraded to K3 Summit (Gatan®) using a slit width of 20 eV.

For the data collected with K2 camera, movies were taken in the super-resolution mode at a nominal magnification 130,000×, corresponding to a physical pixel size of 1.046 Å, and a defocus range from −1.2 to −2.2 μm. Each movie stack was dose-fractionated to 36 frames with a total exposure dose of about 53 $e^-/Å^2$ and exposure time of 7.2 s.

For data collected with K3 camera, movies were taken in the super-resolution mode at a nominal magnification 81,000×, corresponding to a physical pixel size of 1.064 Å, and a defocus range from −1.2 to −2.2 μm. Each movie stack was dose-fractionated to 40 frames with a total exposure dose of about 58 $e^-/Å^2$ and exposure time of 3.0 s.

**Cryo-EM image processing**

The routine processing of all datasets was carried out with the same procedure. Movie stacks were binned 2 × 2, dose weighted, and motion corrected using MotionCor2(*3*) within RELION(*4*). Parameters of contrast transfer function (CTF) were estimated by using Gctf(*5*). Bad images were excluded upon ice condition, defocus range and estimated resolution. Remaining good images were imported into cryoSPARC(*6*) for further patched CTF-estimating, blob-picking and 2D classification. Several good 2D classes were selected as the template for template-picking. From the 2D classification classes, good particles from blob-picking and template-picking were merged and deduplicated. Two rounds of 3D classification were done in RELION, using an initial model generated by cryoSPARC as reference. The particles from the high quality classes were selected and re-extracted without binning. Auto-refinement was performed with C1 symmetry, with further CTF-refinement and particle polishing, yielding consensus maps at the range of 3.4-3.8 Å. Then the well-aligned particles were imported back into cryoSPARC for 3D variability analysis (3DVA) regarding conformational change. Surprisingly, these samples containing two anions all had different conformations including inward-open state and intermediate state. Preliminary data processing was finished so far, and variant strategies were employed to classify different conformations (fig. S5 to S9).

The reported resolutions are all based on the gold-standard Fourier shell correlation (FSC) 0.143 criterion. All the visualization and evaluation of 3D density maps were performed with UCSF Chimera(*7*). The above procedures of data processing are summarized in fig. S5-S9. These sharpened maps were generated by DeepEMhancer(*8*) and then "vop zflip" to get the correct handedness in UCSF Chimera for subsequent model building and structural analysis.

**Model building and structure refinement**

Model building of inward-open state with C2 symmetry was refined from predicted model in Alpha Fold with Coot(*9*) based on the high-resolution 3.3 Å pendrin-Cl cryo-EM map. Most residues were clearly resolved in our cryo-EM map, and totally 665 (18-737Δ596-650) amino acid residues were constructed for each monomer. N-terminal 1-17, IVS  596-650 and C-terminal 738-780 were not modeled because the corresponding density was absent in the map. Structure refinement was performed with PHENIX(*10*) with secondary structure and geometry restraints to prevent structure overfitting. Model building of intermediate state with C2 symmetry was refined based on the 3.9 Å pendrin-Cl/$HCO_3$ cryo-EM map with the same workflow above. Other C1 symmetrical models were refined from C2 models based on their own maps. Statistics associated with data collection, 3D reconstruction and model refinement can be found in table S4.

**Electrophysiological recording**

The whole-cell voltage-clamp recording was carried out in HEK293T cells using an EPC-10 USB amplifier with Patchmaster v2.90.5 software (HEKA Elektronik GmbH, Reutlingen, Germany). The recording micropipettes were pulled from capillary glass (BF150-86-10, Sutter Instrument, Novato, CA, USA) followed by fire-polishing (3-5 MΩ). To analyze chloride transport by pendrin, HEK293T cells were bathed in extracellular solution composed of 146 mM NaCl, 2 mM $MgCl_2$, 5 mM EGTA and 10 mM HEPES (pH 7.4 adjusted with N-methyl-D-glucamine) and in pipette solution containing the same components as the extracellular solution. The holding potential was 0 mV. Currents were acquired by a series of 200 ms depolarizing steps from -100 mV to 100 mV in 20 mV increments. Data were analyzed in Igor Pro 6.22A and GraphPad Prism 8. Statistical significance was evaluated by unpaired t test. Data are shown as means ± SEM.

**Fluorescence anion exchange assay**

Fluorescence assays were taken 48 hours after transfection. Measurements of intracellular pH in HEK 293T cells transiently transfected with mCherry-pendrin plasmids were performed using a pH sensitive fluorescent probe BCECF(2',7'-bis-(2-carboxyethyl)-5-(and-6)-carboxyfluorescein)(*11*). After dye loading (15 min, 37 °C), the cells were perfused with a buffer containing 110 mM NaCl, 25 mM NaHCO3, 10 mM glycose and 5mM HEPES pH 7.5, and BCECF fluorescence was recorded at the excitation wavelengths of 488 nm and the emission wavelengths of 530±10 nm by confocal microscope (Leica TCS SP8) at 37 °C with a 5% CO2 gassing. Then cells were treated alternately by $Cl^-$-free buffer (NaCl is replaced by sodium gluconate) and $Cl^-$-containing buffer and photographed under the same conditions. $Cl^-/HCO_3^-$ exchange activities were estimated from fluorescence intensity change.

For in vivo $Cl^-/I^-$ measurements, HEK 293T were transiently co-transfected with EYFP and mCherry-pendrin plasmids. After titration, we selected the $Cl^-/I^-$ concentration tuning the fluorescence intensity within the appropriate range of fluorescence microscopy. Cells were treated alternately by $Cl^-$-containing buffer (140 mM NaCl, 10 mM glycose and 5mM HEPES pH 7.5) and $I^-$-containing buffer (25 mM NaI, 115 mM sodium gluconate, 10 mM glycose and 5mM HEPES pH 7.5). The EYFP fluorescence was recorded at the excitation wavelengths of 488 nm and the emission wavelengths of 530±10 nm by the same confocal microscope at 37 °C with a 5% CO2 gassing. $Cl^-/I^-$ exchange activities were estimated from fluorescence intensity change(*12*).

The fluorescence intensity was measured by ImageJ and calculated by GraphPad Prism 8.

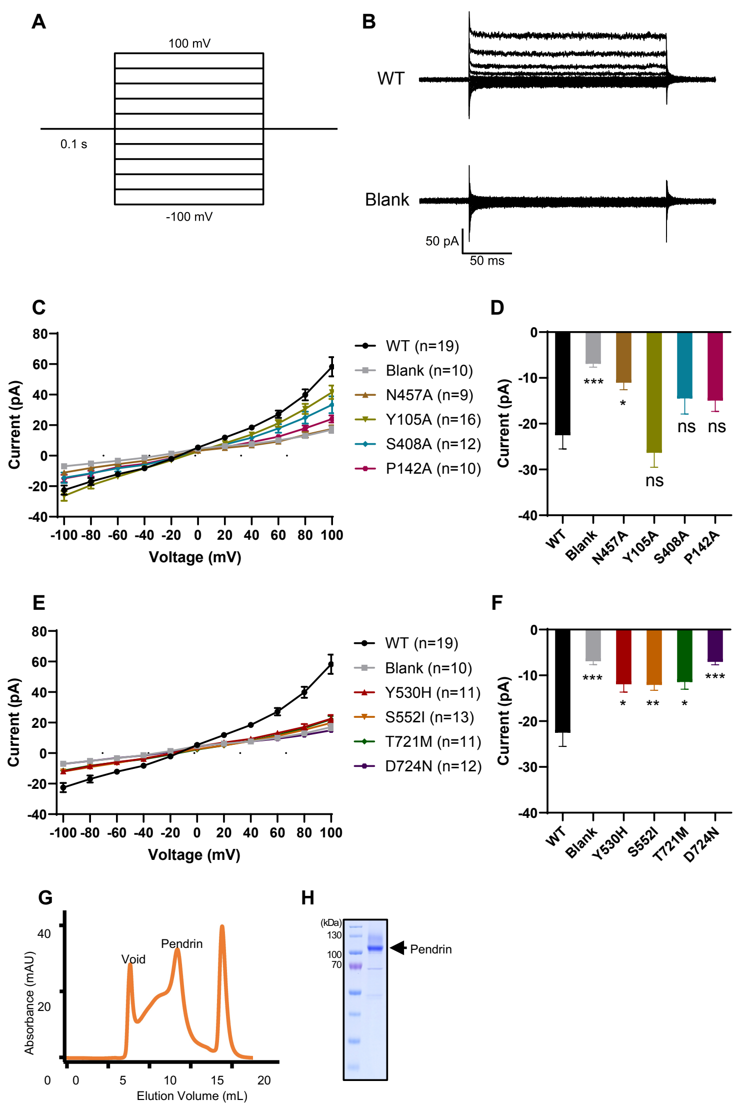


**Fig. S1. Electrophysiology and biochemistry of pendrin.**

(**A**) The stimulation protocol of whole-cell voltage-clamp recording. The experiments were carried out in HEK293T cells with a holding potential of 0 mV. Obtain the currents by a series of 200 ms depolarizing steps between -100 mV and 100 mV in 20 mV increments. (**B**) The representative $Cl^-$ current traces of WT pendrin (up) and Blank (bottom). Blank, non-transfected HEK293T cells. (**C**) The current-voltage relation curves of anion-binding site mutants. The number of collected cells was shown. (**D**) Currents of anion-binding site mutants at the voltage of -100 mV. ***P < 0.001 and *P < 0.05 versus the cells transfected with WT pendrin, unpaired t test. (**E**) The current-voltage relation curves of STAS domain mutants. Chloride currents were recorded. The number of collected cells was shown. (**F**) Chloride currents of STAS domain mutants at the voltage of -100 mV. ***P < 0.001, **P < 0.01 and *P < 0.05 versus the cells transfected with WT pendrin, unpaired t test. (**G**) Size exclusion chromatography of pendrin-$Cl^-$. (**H**) SDS-PAGE gel of the pulled peak fraction of pendrin-$Cl^-$.

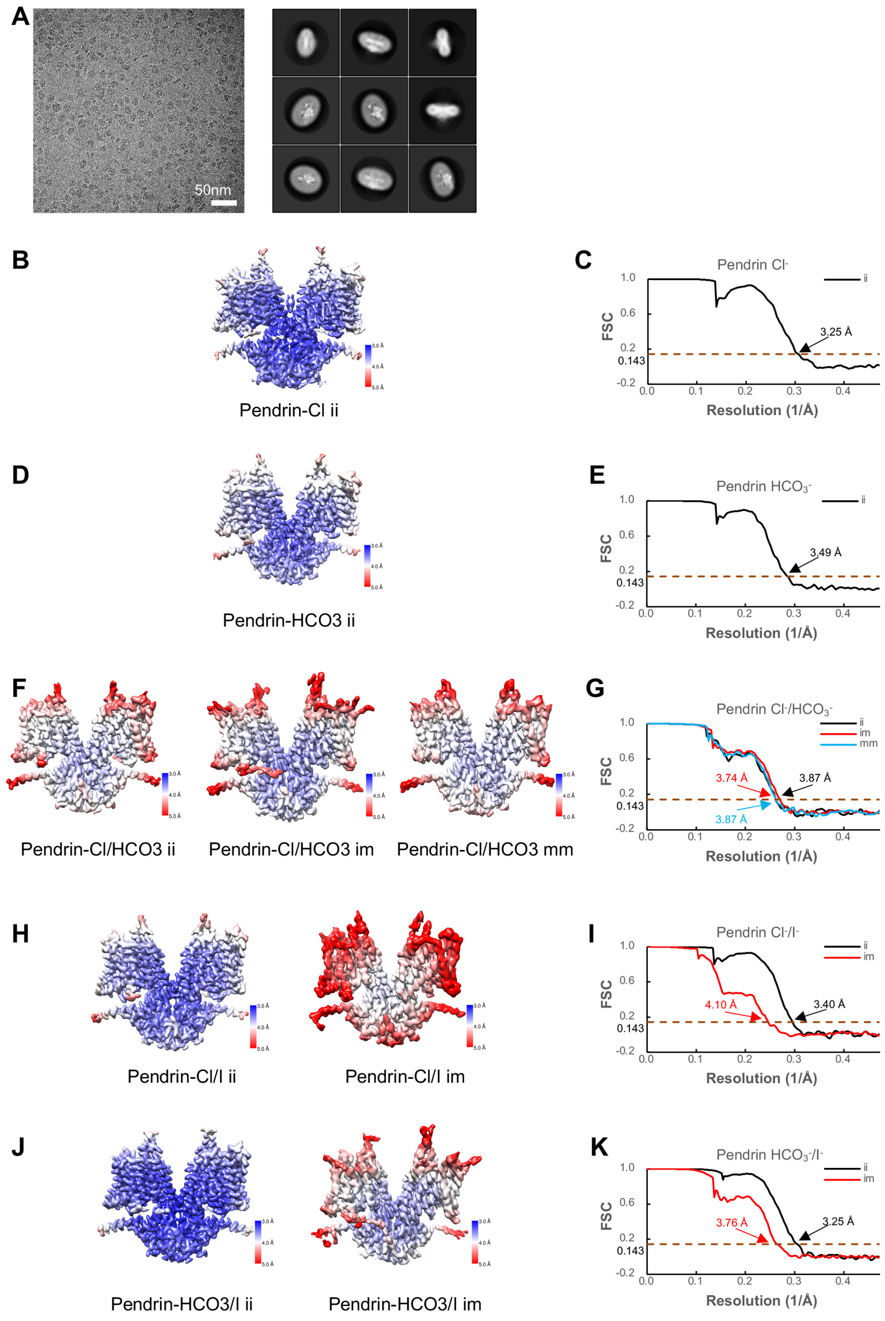


**Fig. S2. Cryo-EM micrograph and data processing results.**

(**A**) Cryo-EM micrograph and 2D average of pendrin-$Cl^-$. (**B, D, F, H, J**) cryo-EM maps colored according to local resolution estimations. (**C, E, G, I, K**) Corresponding FSC curves of the masked cryo-EM maps. ii: inward-inward, im: inward-intermediate, mm: intermediate-intermediate for protomer A and protomer B, respectively.

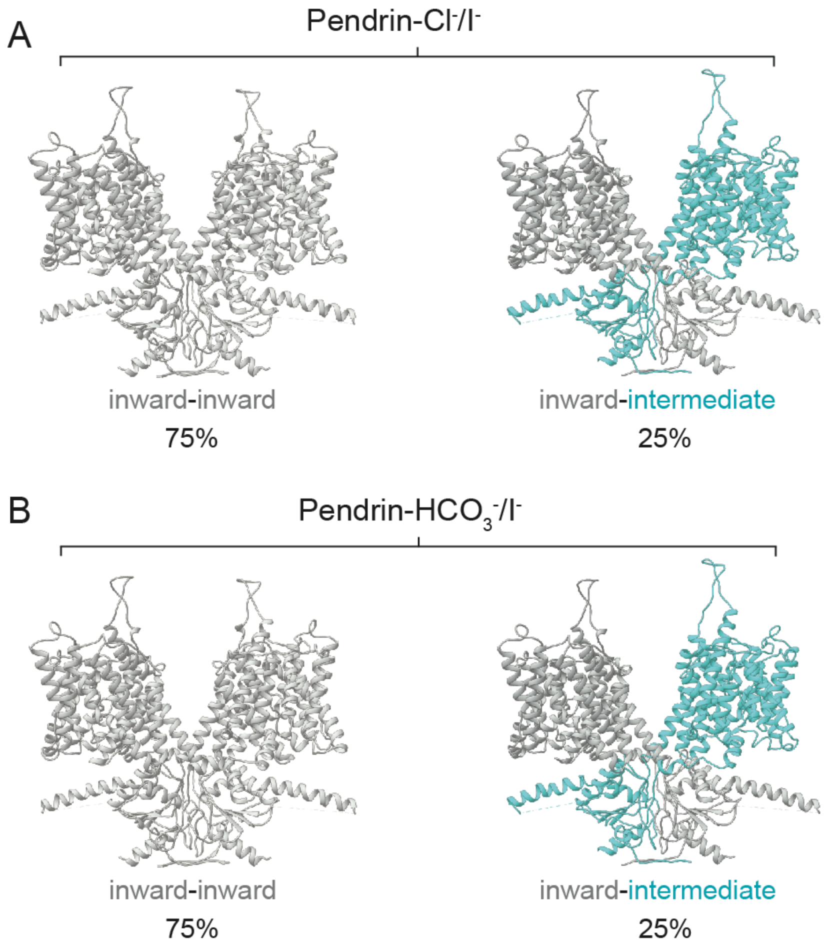


**Fig. S3. Conformations of pendrin-$Cl^-/I^-$ and $HCO_3^-/I^-$.**

(**A**) Two conformations of pendrin-$Cl^-/I^-$. (**B**) Two conformations of pendrin-$HCO_3^-/I^-$. Proportion of particles is indicated.

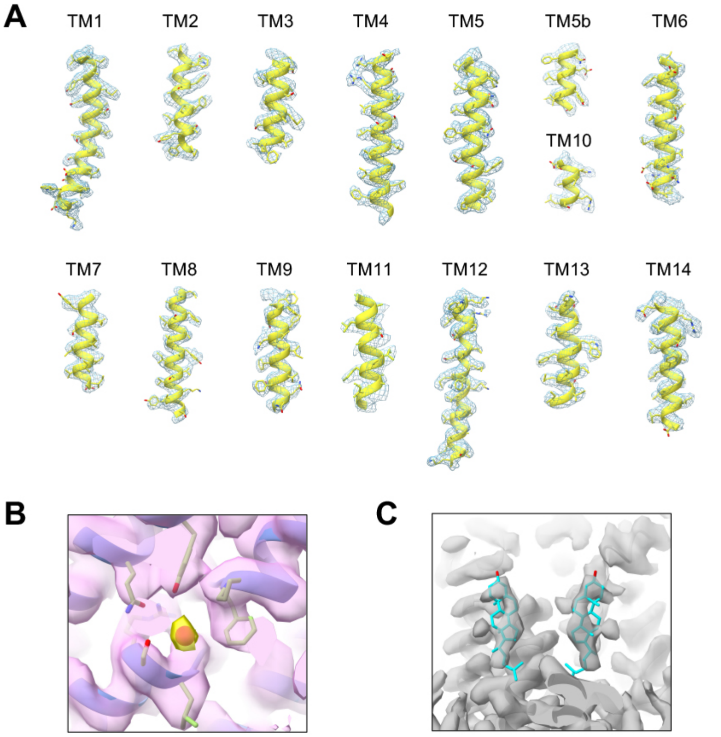


**Fig. S4. Local cryo-EM densities of pendrin-Cl$^-$ structures.**

(**A**) Representative local cryo-EM densities of pendrin-Cl$^-$. (**B**) Cryo-EM densities near the anion binding site in pendrin-Cl$^-$. Density representing Cl$^-$ is colored in yellow. (**C**) Cryo-EM densities representing cholesterol. Cholesterol is shown in cyan stick representation.

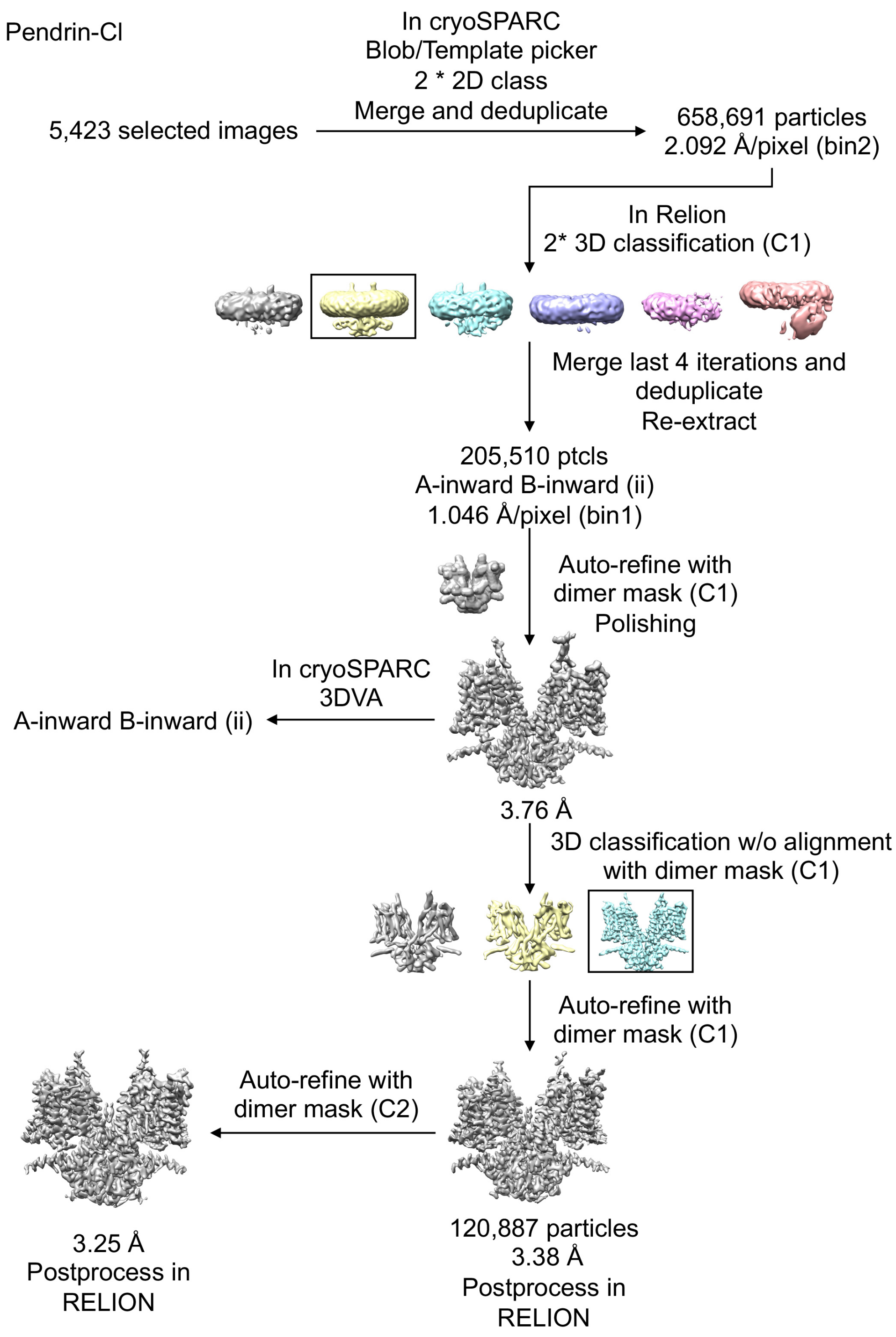


**Fig. S5. Flowchart of pendrin-Cl$^-$ data processing**

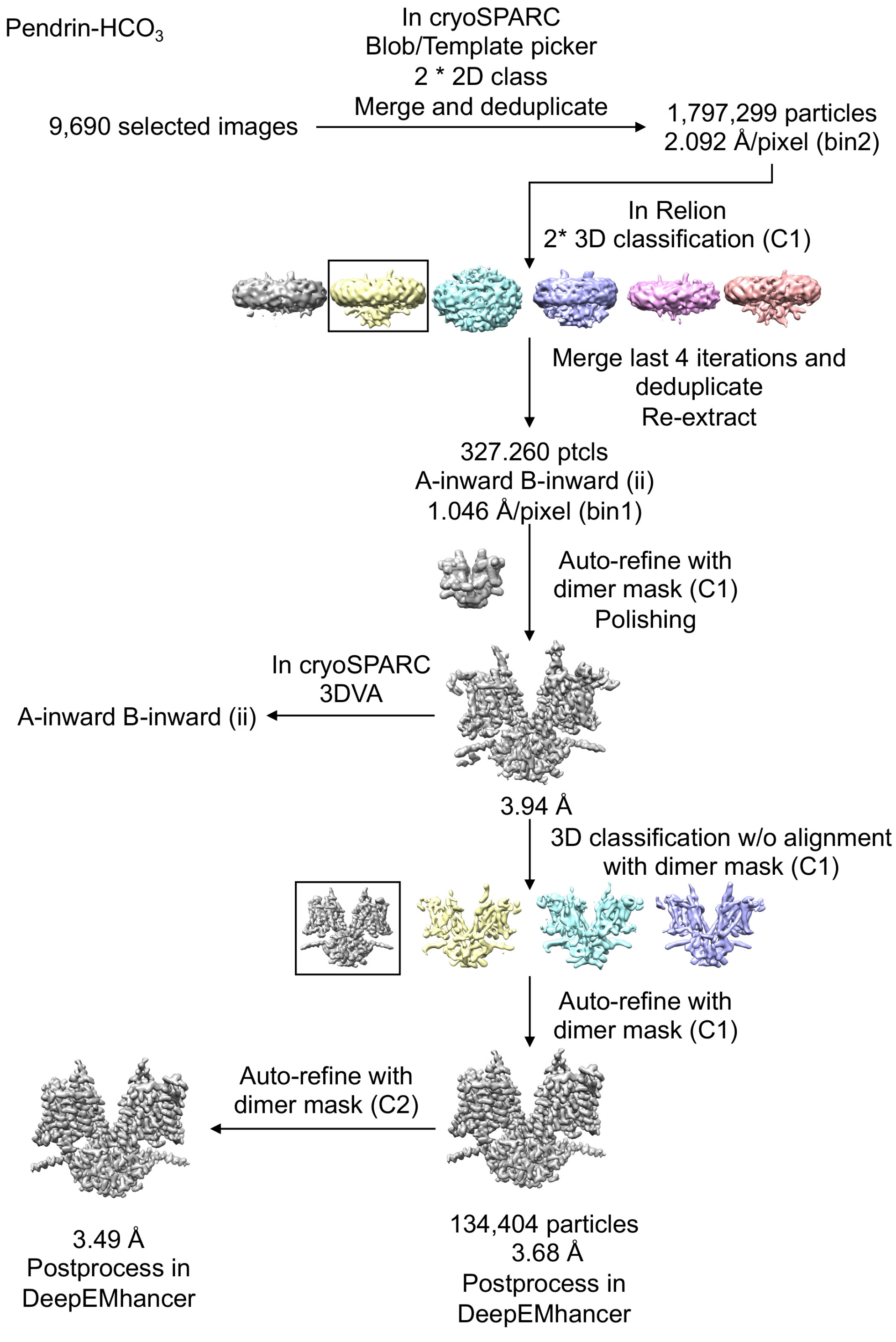


**Fig. S6. Flowchart of pendrin-$HCO_3^-$ data processing**

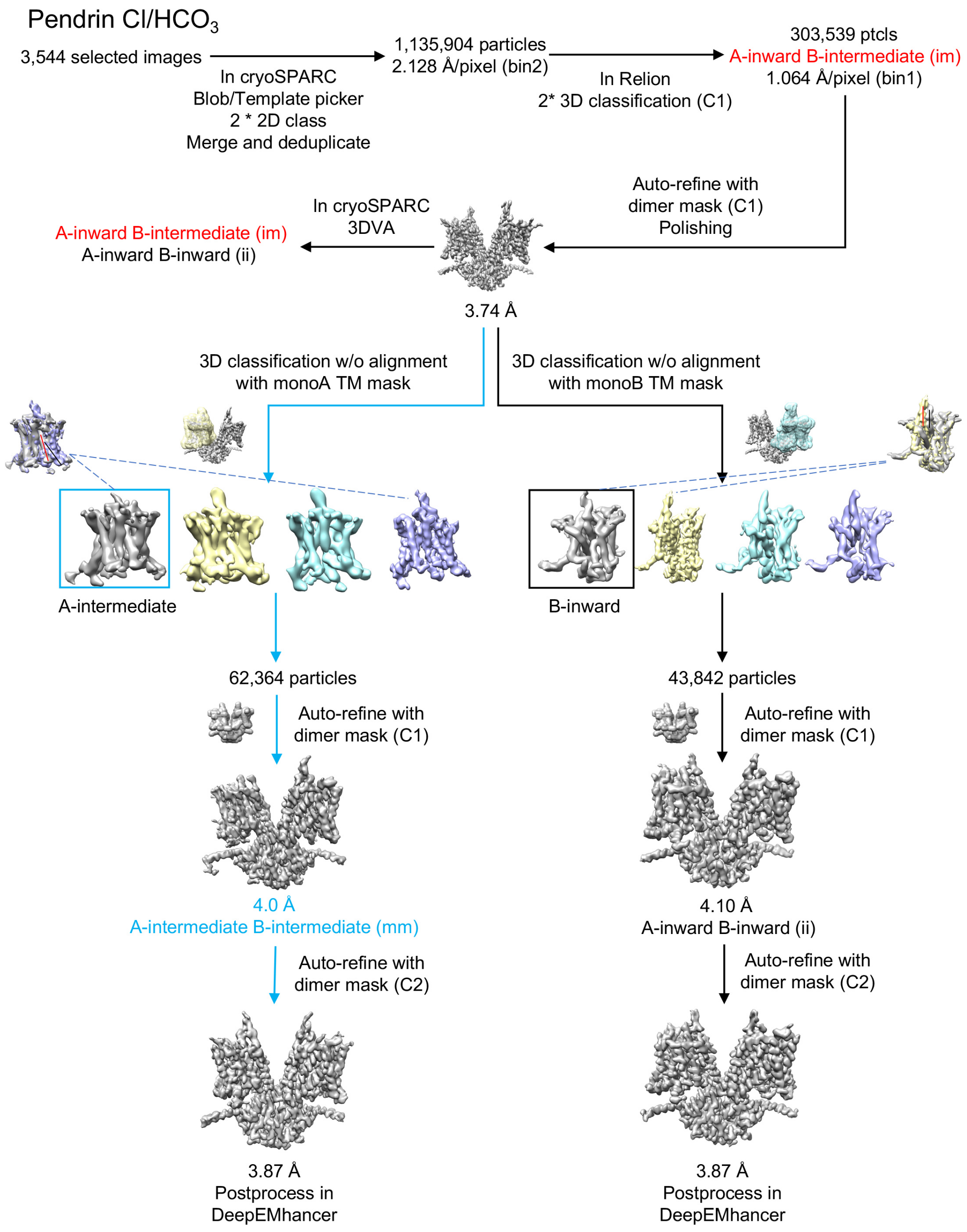


**Fig. S7. Flowchart of pendrin-$Cl^-/HCO_3^-$ data processing**

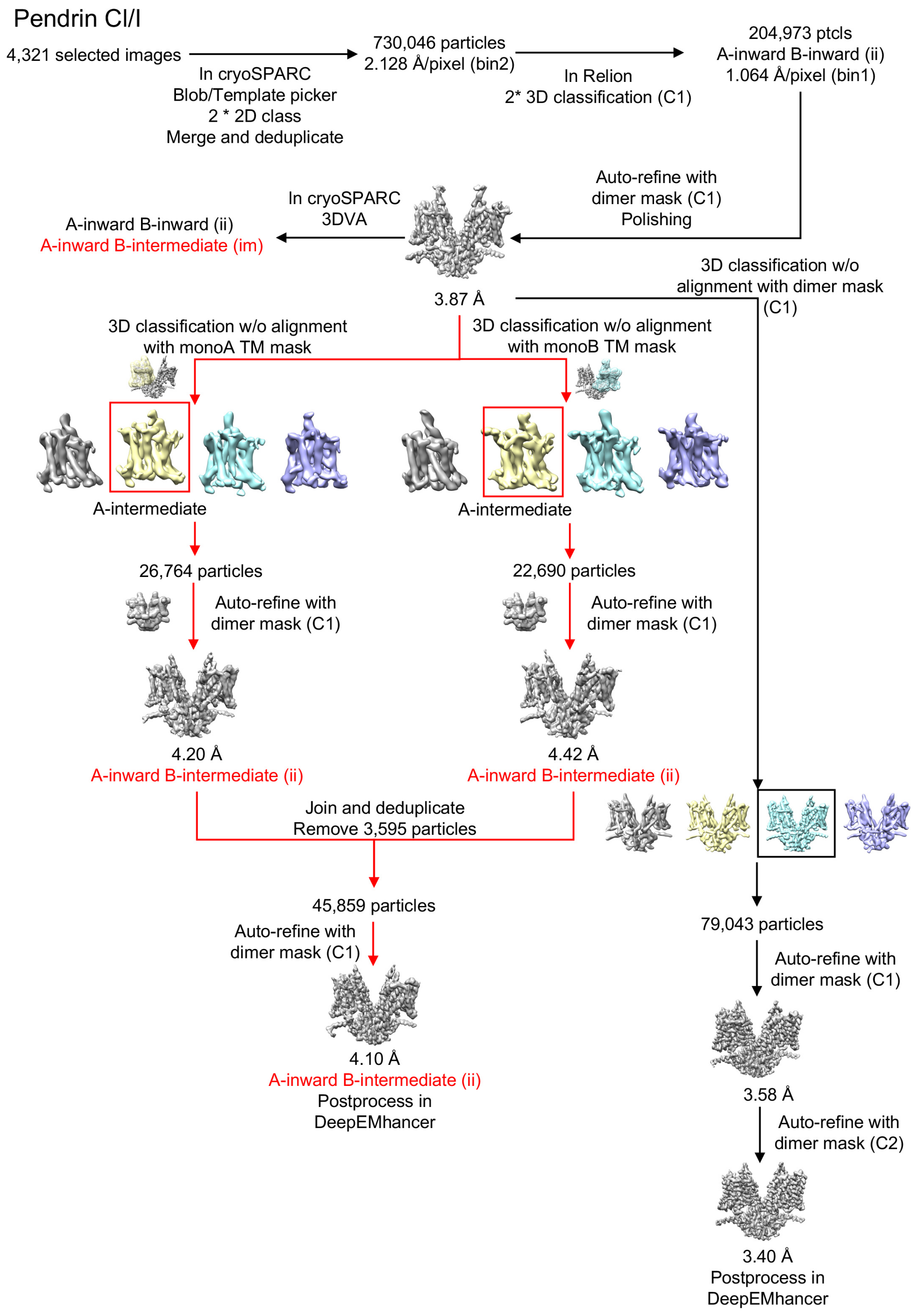


**Fig. S8. Flowchart of pendrin-Cl$^-$/I$^-$ data processing**

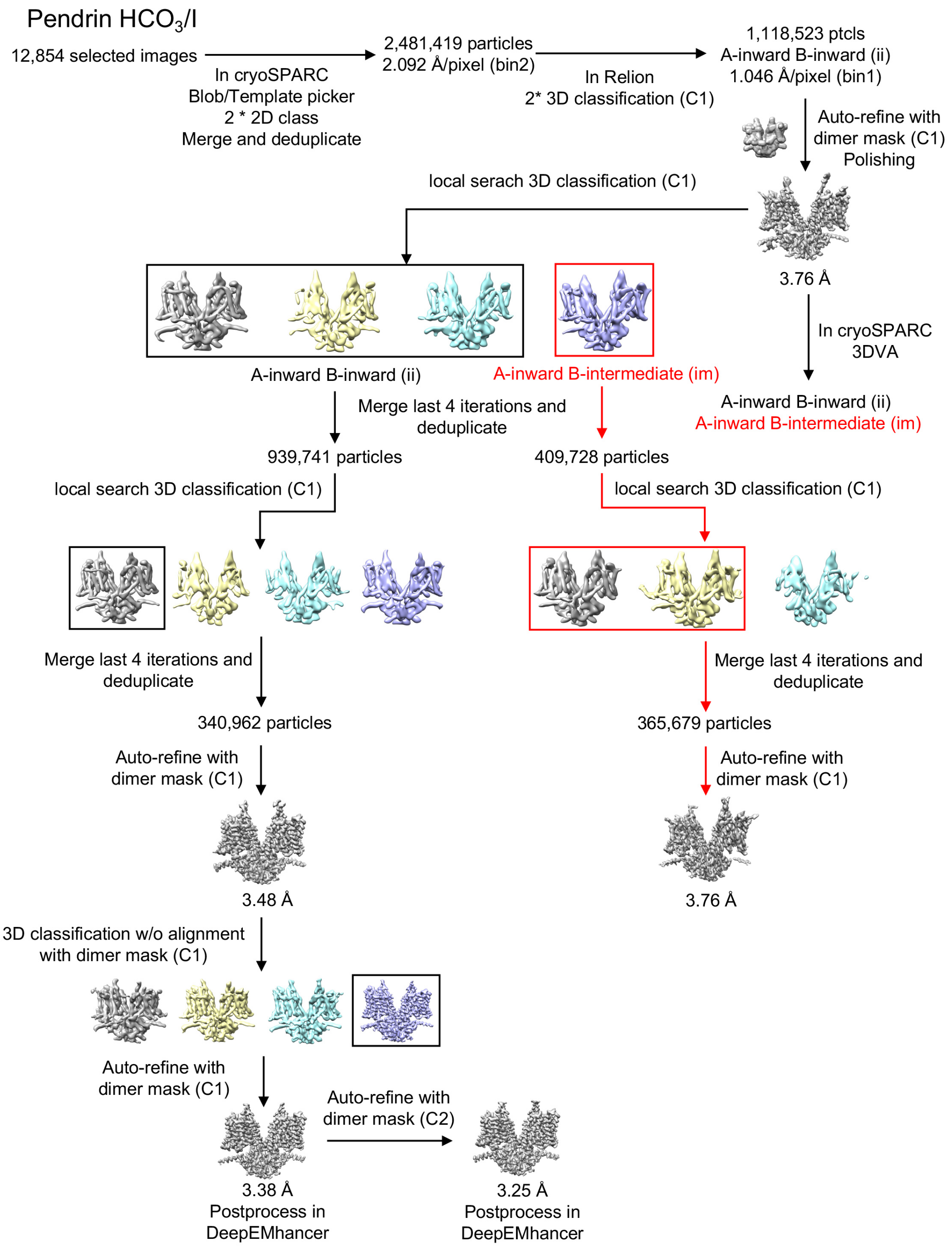


**Fig. S9. Flowchart of pendrin-$HCO_3^-$/$I^-$data processing**

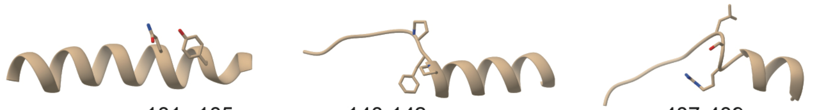

```
                                     101  105          140-142                 407-409
Q9R155|S26A4_MOUSE    ISGVSTGLVGTLQGMAYALL  HISVGPFPVVSLMVGSVVLS  FVATTALSRTAVQESTG
O43511|S26A4_HUMAN    ISGVSTGLVATLQGMAYALL  HISVGPFPVVSLMVGSVVLS  FVATTALSRTAVQESTG
Q8BU91|S26A9_MOUSE    LGGLSGGCIQVPQGMAFALL  QMVPGTFAVISILVGNICLQ  HVICCALSVTLAVDGAG
Q7LBE3|S26A9_HUMAN    LGGLSGGSIQVPQGMAFALL  QMVPGTFAVISILVGNICLQ  HVICCALSVTLAVDGAG
Q99NH7|S26A5_MOUSE    VSGISTGVLQLPQGLAFAML  HISIGPFAVISLMIGGVAVR  FSISCSLSRSLVQEGTG
P58743|S26A5_HUMAN    VSGISTGVLQLPQGLAFAML  HISIGPFAVISLMIGGVAVR  FSISCSLSRSLVQEGTG
Q9EPH0|S26A5_RAT      VSGISTGVLQLPQGLAFAML  HISIGPFAVISLMIGGVAVR  FSISCSLSRSLVQEGTG
D7PC76|D7PC76_TURTR   VSGISTGVLQLPQGLAFAML  HISIGPFAVISLMIGGVAVR  FAISCSLSRSLVQEGTG
                      :.*:* * :   **:*:*:*  ::   * * *:*:::*.: :  .    :** : . :.:*
```

**Fig. S10. SLC26A homologues sequence alignment around the binding pocket.**

These sequences information can be found on Uniport. The entry identifier is listed along with the protein name in the left.

**Table S1. Comparison of pendrin structures at different conditions (R.m.s.d).**

| | | | | | | | | | | | | |
|---|---|---|---|---|---|---|---|---|---|---|---|---|
| pendrin-Cl | | | | | | | | | | | | |
| Pendrin-$HCO_3$ | 0.3 (Å) | | | | | | | | | | | |
| pendrin-Cl/$HCO_3$ ii | 0.4 | 0.4 | | | | | | | | | | |
| pendrin-Cl/I ii | 0.3 | 0.4 | 0.3 | | | | | | | | | |
| pendrin-$HCO_3$/I ii | 0.3 | 0.4 | 0.4 | 0.4 | | | | | | | | |
| pendrin-Cl/$HCO_3$ im A | 0.5 | 0.5 | 0.4 | 0.5 | 0.5 | | | | | | | |
| pendrin-Cl/$HCO_3$ im B | 4.4 | 4.6 | 4.3 | 4.3 | 4.3 | 4.3 | | | | | | |
| pendrin-Cl/I im A | 0.7 | 0.7 | 0.6 | 0.6 | 0.6 | 0.6 | 4.1 | | | | | |
| pendrin-Cl/I im B | 4.1 | 4.2 | 4.0 | 4.1 | 4.1 | 4.0 | 0.8 | 3.9 | | | | |
| pendrin-$HCO_3$/I im A | 0.7 | 0.7 | 0.7 | 0.7 | 0.7 | 0.6 | 4.4 | 0.8 | 4.2 | | | |
| pendrin-$HCO_3$/I im B | 4.3 | 4.4 | 4.2 | 4.2 | 4.3 | 4.2 | 0.6 | 4.1 | 0.9 | 4.3 | | |
| pendrin-Cl/$HCO_3$ mm | 4.5 | 4.6 | 4.4 | 4.4 | 4.4 | 4.4 | 0.6 | 4.2 | 0.9 | 4.5 | 0.8 | |
| | pendrin-Cl | pendrin-$HCO_3$ | pendrin-Cl/$HCO_3$ ii | pendrin-Cl/I ii | pendrin-$HCO_3$/I ii | pendrin-Cl/$HCO_3$ im A | pendrin-Cl/$HCO_3$ im B | pendrin-Cl/I im A | pendrin-Cl/I im B | pendrin-$HCO_3$/I im A | pendrin-$HCO_3$/I im B | pendrin-Cl/$HCO_3$ mm |

**Table S2. Comparison of pendrin structures with representative structures of prestin and SLC26A9 (R.m.s.d).**

| | | | | | | | | | | |
|---|---|---|---|---|---|---|---|---|---|---|
| Pendrin-Cl | | | | | | | | | | |
| SLC26A9 (6RTC) | 4.6 (Å) | | | | | | | | | |
| SLC26A9 (7CH1) | 4.1 | 3.4 | | | | | | | | |
| Prestin (7S9C) | 3 | 5.3 | 5.3 | | | | | | | |
| Prestin (7LH2) | 3.8 | 6.3 | 6.3 | 2.3 | | | | | | |
| Prestin (7S9D) | 3.1 | 6.1 | 6 | 2.1 | 1.9 | | | | | |
| Prestin (7S8X) | 3.6 | 6.9 | 7.2 | 2.7 | 1.6 | 1.9 | | | | |
| Prestin (7LGU) | 4.5 | 7.5 | 8 | 3.1 | 1.2 | 2.4 | 1.4 | | | |
| SLC26A9 (6RTF) | 4.3 | 3 | 4.8 | 4.3 | 4.4 | 4.8 | 5.1 | 5.8 | | |
| Pendrin-Cl/$HCO_{3\ mm}$ | 4.5 | 7 | 6.8 | 4.4 | 4.2 | 3.8 | 3.3 | 3.9 | 5.1 | |
| | Pendrin-Cl | SLC26A9(6RTC) | SLC26A9(7CH1) | Prestin (7S9C) | Prestin (7LH2) | Prestin (7S9D) | Prestin (7S8X) | Prestin (7LGU) | SLC26A9(6RTF)) | Pendrin-Cl/$HCO_{3\ mm}$ |

### Table S3. Pathogenetic missense variants.

| Residue change | Protein location | Cellular localization | Functionality | Predicted molecular effect |
|---|---|---|---|---|
| S28R | NTD-Nα1 | PM | Loss of Cl- uptake | Disruption of H bond with E29; interfere the relative folding and orientation surround |
| G102R | Core-TM1 | ER | | Clash between TM1 and TM2; Insertion of charged residue in a hydrophobic interface between TM1 and TM2 |
| P123S | Core-TM2 | Intracell. | | Disruption of hydrophobic interaction between TM9 and TM10; destabilize the binding pocket |
| V138F | Core-TM3 loop | ER | | Destabilize the binding pocket |
| R185T | Core-TM4 | Intracell. | | Change the positive charge of surface |
| G209V | Intracellular loop between Core and Gate | PM | Severe reduction of I- transport | Change the interface between TMD and STAS |
| L236P | Gate-TM5 | ER, Intracell. | | Insertion of Pro in α-helix; introduce a kink and leads to the misfolding of TM5 |
| V239D | Gate-TM5 | ER | | Disruption of hydrophobic interaction between TM5 and TM6 |
| E303Q | Gate-TM7 | PM | Loss of Cl-/ I- and Cl-/HCO3- exchange activity | Change the positive charge of surface |
| F335L | Hydrophobic loop between TM7 and TM8 | PM | Reduction of Cl-/ I- and Cl-/HCO3- exchange activity | Disruption of hydrophobic interaction with lipids |
| E384G | Core-TM9 | ER, Intracell. | | Disruption of H bond with Y127 necessary for TM9 and TM2 interaction |
| N392Y | Core-TM9 | Intracell. | | Larger side chain; clash between TM9 and TM10; destabilize the binding pocket |
| R409H | Core-TM10 | Partially PM | Reduction of Cl- and I- transport | Destabilize the binding pocket |
| T410M | Core-TM10 | ER | | Larger side chain; destabilize the binding pocket |
| L445W | Loop between Core and Gate | ER, Intracell. | | Larger side chain, change in steric hindrance |
| G497S | Gate-TM14 | Intracell. | | Disruption of interaction between TM13 and TM14 |
| Y530H/S | STAS-Loop | ER, Intracell. | | Interfere local folding of the long loop |
| Y556C | STAS-Loop | Partially PM | Loss of I- transport | Disruption of hypothetical ion binding pocket |
| C565Y | STAS-Cα1 | PM | Reduction of Cl-/ I- and Cl-/HCO3- exchange activity | Larger side chain, change in steric hindrance |
| G672E | STAS-Cα2 | Partially PM | Loss of I- transport | Disruption of hypothetical ion binding pocket |
| T721M | STAS-Cα4 | Intracell. | | Disruption of H bond with D724; affect the folding of Cα4 and decrease the solubility of STAS domain. |
| H723R | STAS-Cα4 | ER, Intracell. | | Affect hypothetical interactions surround; interfere the relative folding and orientation of Cα4. |

### Table S4. Cryo-EM data collection and refinement statistics.

| | pendrin-Cl | pendrin-$HCO_3$ | pendrin-Cl/I ii | pendrin-Cl/I im | pendrin-Cl/$HCO_3$ ii | pendrin-Cl/$HCO_3$ im | pendrin-Cl/$HCO_3$ mm | pendrin-$HCO_3$/I ii | pendrin-$HCO_3$/I im、 |
|---|---|---|---|---|---|---|---|---|---|
| **PDB ID** | 7WK1 | 7WK7 | 7WL8 | 7WLB | 7WL7 | 7WL9 | 7WLE | 7WL2 | 7WLA |
| **EMDB** | 32555 | 32561 | 32577 | 32580 | 32576 | 32578 | 32583 | 32574 | 32579 |
| **Data collection and processing** | | | | | | | | | |
| Data sets | pendrin-Cl | pendrin-$HCO_3$ | pendrin-Cl/I | | pendrin-Cl/$HCO_3$ | | | pendrin- $HCO_3$/I | |
| Voltage (kV) | 300 | 300 | 300 | | 300 | | | 300 | |
| Detector | K2 | K2 | K3 | | K3 | | | K2 | |
| Pixel size (Å) | 1.046 | 1.046 | 1.064 | | 1.064 | | | 1.046 | |
| Electron dose ($e^-/Å^2$) | 53 | 53 | 58 | | 58 | | | 53 | |
| Defocus range (μm) | −1.2 to −2.2 | −1.2 to −2.2 | −1.2 to −2.2 | | −1.2 to −2.2 | | | −1.2 to −2.2 | |
| Final particles | 120,887 | 134,404 | 79,043 | 45,859 | 43,842 | 197,333 | 62,364 | 221,446 | 365,679 |
| Final resolution (Å) | 3.25 | 3.49 | 3.40 | 4.10 | 3.87 | 3.74 | 3.87 | 3.25 | 3.76 |
| **Model refinement and validation statistics** | | | | | | | | | |
| Ramachandran statistics | | | | | | | | | |
| Favored (%) | 94.70 | 94.93 | 95.31 | 95.45 | 95.31 | 95.30 | 93.92 | 95.01 | 94.92 |
| Allowed (%) | 5.22 | 4.99 | 4.61 | 4.55 | 4.61 | 4.62 | 6.08 | 4.92 | 5.08 |
| Outliers (%) | 0.08 | 0.08 | 0.08 | 0.00 | 0.08 | 0.08 | 0.00 | 0.08 | 0.00 |
| Rotamer outliers (%) | 3.12 | 2.94 | 1.87 | 3.92 | 3.39 | 4.27 | 5.25 | 2.14 | 4.01 |
| R.m.s.deviations | | | | | | | | | |
| Bond lengths (Å) | 0.002 | 0.001 | 0.002 | 0.002 | 0.001 | 0.002 | 0.002 | 0.002 | 0.002 |
| Bond angles (º) | 0.406 | 0.406 | 0.402 | 0.400 | 0.409 | 0.403 | 0.426 | 0.412 | 0.405 |
